# Genetic Modulation of Oxycodone Self-Administration Trajectories: From Initiation to Escalating Burst Patterns

**DOI:** 10.64898/2026.06.15.732499

**Authors:** Caleb I. Hodges, Eamonn P. Duffy, Jonathan O. Ward, Luanne H. Hale, Cove Andrews, Laura M. Saba, Marissa A. Ehringer, Ryan K. Bachtell

**Affiliations:** Department of Psychology and Neuroscience, University of Colorado Boulder, Boulder, CO, United States; Department of Integrative Physiology, University of Colorado Boulder, Boulder, CO, United States; Institute for Behavioral Genetics, University of Colorado Boulder, Boulder, CO, United States; Department of Pharmaceutical Sciences, Skaggs School of Pharmacy and Pharmaceutical Sciences, University of Colorado Anschutz Medical Campus, Aurora, CO, United States

**Author notes:** **Corresponding Author:** Ryan K. Bachtell.

**Keywords:** Inbred, Drug Loading, Compulsive, Inter-Infusion Interval, Addiction, Substance Abuse, Sex-differences, Intermittent, Long Access, Short Access

## Abstract

Opioid Use Disorder (OUD) remains a prominent threat to global health. Genetic background influences the susceptibility of developing OUD, although specific genetic factors remain elusive. Rodent models that differ in susceptibility to escalation and dysregulation of opioid use are valuable tools to facilitate discovery of genetic pathways. Phenotypes associated with the development of OUD were compared in seven classic inbred rat strains (M520/N, WKY/NCrl, F344/NCrl, F344/Stm, LEW/Crl, LEW/SSNHsd, LE/Stm) from the Hybrid Rat Diversity Panel (HRDP). A two-phase self-administration paradigm was utilized to assess characteristics of the acquisition of oxycodone self-administration during daily 2-h sessions, and the escalation of oxycodone use during daily 12-h sessions. Genetic background influenced the acquisition of oxycodone self-administration as indicated by differences in the initiation of responding for oxycodone during each session and different amounts of oxycodone intake. We observed that escalation of oxycodone intake between-sessions was strain dependent, and the within-session distribution of oxycodone intake was strongly influenced by strain. The M520/N strain engaged in a unique pattern of intake, characterized by rapid initiation of oxycodone responding during the acquisition phase and a significant burst-like responding during escalation. Strain-dependent sex differences were also observed in several acquisition and escalation metrics. Of interest, burst responding was more prevalent in females of the M520/N strain compared to males. Together, these data indicate that genetic background influences not only overall oxycodone intake, but specific within- and between-session metrics that capture patterns of consumption across the substance use trajectory.

## 1. Introduction

Despite being one of the most prescribed drug classes in the United States, opioids are long recognized for their abuse potential (1). Prescription of opioids including oxycodone has been associated with the start of the opioid epidemic, a crisis that has resulted in approximately 6.7 to 7.6 million individuals in the United States developing opioid use disorder (OUD, 2). In response to the opioid epidemic, opioid prescriptions have steadily declined since 2011 (3). However, there is evidence suggesting that prescription opioids remain effective in certain populations where alternative treatments are lacking, such as in cancer patients (4,5). Opioids are highly effective at producing acute analgesia, but long-term use of opioids can result in dependence, tolerance, and overdose (6–8). Understanding which individuals in these populations are vulnerable or resistant to developing OUD will aid clinical decisions regarding pain treatment.

Although there is high abuse potential associated with opioids, OUD only occurs in a subset of individuals who regularly consume opioids (9). Nearly 13% of people who used prescription opioids have misused opioids with approximately 16% of those individuals developing OUD (10,11). Heritability estimates of OUD range from 0.31-0.54, suggesting that individuals differ in their genetic susceptibility to developing problematic opioid use (12–14). Genome-wide association studies (GWAS) have identified few genetic variants associated with the development of OUD (15). These datasets often rely on broad diagnostic criteria, which limits their ability to identify genetic variants associated with protection or vulnerability across different phases of drug use such as initiation, maintenance of drug use, or compulsive use that is observed in OUD (16). Animal models can provide insight into the genetic contribution by examining specific behavioral phenotypes relevant to the progression of OUD that cannot be readily assessed in human cross-sectional GWAS. For example, a recent GWAS conducted in heterogenous stock (HS) rats revealed that heroin consumption and extinction are heritable traits, and identified genetic variants associated with heroin consumption, escalation of intake, and motivation to obtain heroin (17).

Phenotypic characterization of opioid-related behaviors in genetically diverse rodent populations provides a powerful approach for identifying genetic factors that contribute to OUD. The Hybrid Rat Diversity Panel (HRDP) consists of nearly 100 genetically divergent inbred rat strains, including two recombinant inbred panels (HXB/BXH and FXLE/LEXF) as well as several classic inbred strains (18). Genetic variation across HRDP strains makes it useful for investigating the genetic contributions to complex behavioral traits such as oxycodone-induced analgesia and self-administration (19–23). Work in HS rats further demonstrates substantial individual variability in oxycodone self-administration, escalation, and addiction-like behaviors (24). Together, these rodent models provide complementary platforms for integrating behavioral phenotyping with genomic analyses to identify biological mechanisms contributing to OUD risk.

Intravenous drug self-administration (IVSA) assesses volitional drug intake in rodents with procedural variations such as short-access (ShA) and long-access (LgA) conditions modeling different phases of substance use disorders (25). For example, ShA conditions often facilitate initiation of drug use or acquisition, and often display stabilized intake while LgA conditions promote escalation and compulsive use (26). Within-session patterns of drug self-administration have become increasingly recognized as an important factor in characterizing drug intake throughout the different phases. Belin et al. (2009) found that within-session distribution of cocaine infusions was positively correlated with addiction severity. Moreover, the emergence of a burst-like pattern of responding, where several infusions are delivered within a short duration, predicted high cocaine use severity (27). Recent studies have observed burst-like patterns of heroin self-administration under specific self-administration parameters where rats have periods of free access to heroin (28,29). Analysis of other within-session parameters, such as inter-infusion intervals, suggests that short bursts of oxycodone intake correlate with the reinstatement of oxycodone seeking (30). Little is known about the genetic factors contributing to within-session regulation of drug intake that may reflect the emergence of dysregulated and uncontrolled oxycodone use.

In this study, we utilized a two-phase IVSA paradigm to characterize oxycodone use in male and female rats from several “classic” inbred strains that we previously characterized (21). The inbred rat strains being assessed include a subset of classic, non-recombinant inbred strains that displayed robust escalation of oxycodone intake under LgA conditions (21). Here, we further describe within-session phenotypes associated with the acquisition (ShA) and escalation (LgA) phases of the paradigm. Acquisition was assessed using between-sessions measures such as infusions per session and lever discrimination while within-session measures include early session metrics that measure the initiation of oxycodone responding during each ShA sessions. Escalation was assessed using the between-session measures of the number of infusions per session and the magnitude and rate of escalation, as well as within-session measures such as inter-infusion intervals and burst responding. Combining these between- and within-session metrics across both phases enables us to identify genetic differences between specific phenotypes across the trajectory of oxycodone use. We hypothesized that we would identify differences between the strains in some of the within-session metrics that were not captured by the between-session metrics. While all strains acquired and escalated oxycodone intake, only the M520/N strain displayed a rapid transition to uncontrolled oxycodone use marked by higher burst responding compared to the other strains. These findings suggest that there are genetic influences that contribute to different phases of opioid use and specific characteristics of opioid use patterns.

## 2. Methods

### 2.1 Subjects

Adult (PND60+) male and female rats (n = 193) from seven inbred strains were tested in this experiment (**Table 1**). The WKY/NCrl, F344/NCrl, and LEW/Crl strains were obtained from Charles River. The LEW/SSNHsd (LEW/Ss) strain was obtained from Envigo/Inotiv. The remaining strains (M520/N, F344/Stm, LE/Stm) were obtained from the Medical College of Wisconsin (provided by Dr. Melinda Dwinell, R24OD024617). Rats were allowed to habituate for at least one week following arrival before beginning any procedures. Prior to surgery and the self-administration paradigm, animals were tested for somatosensory and analgesia-related measures (20). Rats were single-housed in a temperature (22°C) and humidity (40%) controlled vivarium that had a 12-h light and 12-h dark cycle. Standard rat chow (Teklad 2918, Envigo/Inotiv, Indianapolis, IN) and water were provided *ad libitum*. Animals were tested in 12 cohorts (n = 4-24/cohort). All procedures were approved and conducted in accordance with institutional guidelines outlined by the Institutional Animal Care and Use Committee at the University of Colorado Boulder, an Association for Assessment and Accreditation of Laboratory Animal Care International accredited institution.

**Table 1.** Sample size per strain and sex.

| Strain | RRID | Oxycodone |  | Saline |  |
| --- | --- | --- | --- | --- | --- |
|  |  | M | F | M | F |
| M520/N | RRRC_00168 | 8 | 9 | 6 | 7 |
| WKY/NCrl | RGD_1358112 | 8 | 7 | 8 | 8 |
| F344/NCrl | RGD_737926 | 9 | 8 | 8 | 8 |
| F344/Stm | RGD_1302686 | 10 | 5 | 10 | 6 |
| LEW/SSNHsd | RGD_737922 | 5 | 5 | 5 | 4 |
| LEW/Crl | RGD_737932 | 6 | 6 | 6 | 5 |
| LE/Stm | RGD_629485 | 4 | 8 | 7 | 7 |
| **Total** |  | **50** | **48** | **50** | **45** |

### 2.2 Intravenous Catheter Surgeries

Jugular indwelling catheters were implanted utilizing previously described procedures (31). Briefly, isoflurane (2-4%) was used to anesthetize animals. Intravenous catheters were inserted into the right jugular vein under aseptic conditions. After isolation of the vein, a 22-gauge needle was used to puncture the vein and allow for catheter tubing implantation which was secured using suture threads. Sterile catheters (Access Technologies, Skokie, IL) consisted of 14 cm of polyurethane tubing (0.51 mm inner diameter, 1.12 mm outer diameter) secured to a 22-gauge back mount pedestal (Protech International). The analgesic carprofen (5 mg/kg, s.c.) and antibiotic enrofloxacin (5 mg/kg, s.c.) were administered immediately prior to surgery. Daily subcutaneous injections of carprofen (5 mg/kg) were administered for two days following surgery while monitoring animal health. Catheters were flushed daily with 0.2 mL heparinized (20 IU/ml) bacteriostatic saline containing gentamicin sulfate (0.33 mg/ml; Hospira) until self-administration procedures began. Catheter patency was ensured using 0.2 mL propofol (10 mg/mL propofol, Sagent) solution administered before the acquisition phase, prior to the escalation phase, and following the escalation phase to ensure patency of the catheters. Prior to self-administration, rats were assigned to the saline or oxycodone group using a pseudorandomized procedure based on analgesic testing (**Figure 1**; 23).

**Figure 1:**
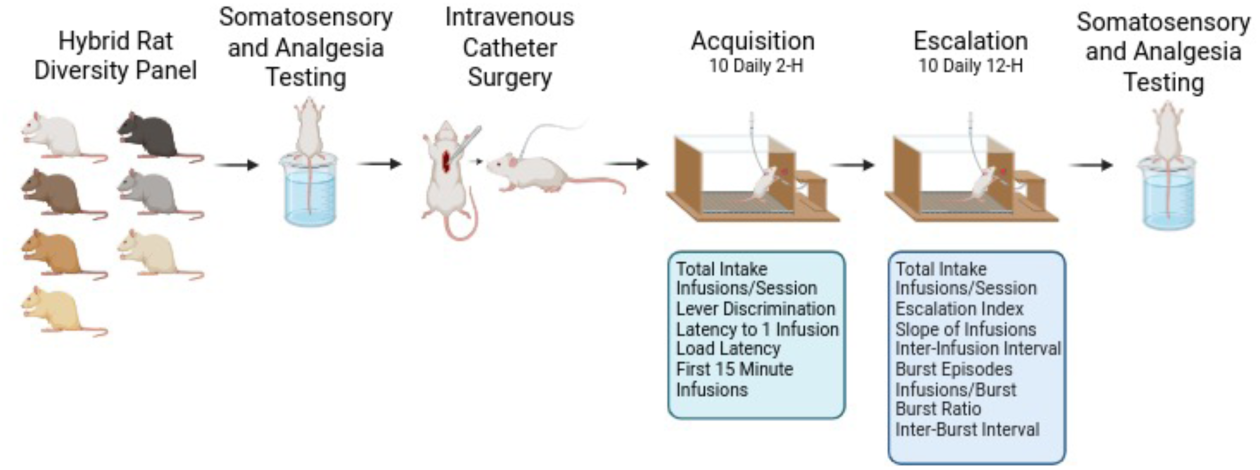
Overview of Longitudinal Behavioral Protocol. Schematic depicting the sequencing of the behavioral phenotyping and surgical procedures. Rats underwent somatosensory and analgesia testing as previously reported before surgical implantation of intravenous catheters (24). Oxycodone or saline intake was evaluated using a self-administration procedure with a fixed-ratio 1 schedule of reinforcement. During the acquisition phase, rats lever pressed for saline/oxycodone in ten daily 2-h (ShA) sessions and behavioral phenotypes were included to evaluate changes in reinforcement learning. The escalation phase was comprised of ten daily 12-h (LgA) sessions and monitored between session changes in intake as well as within-session metrics to assess the temporal organization and dysregulation of drug intake. Rats were subsequently tested in the same somatosensory and analgesia tests (data not shown). Created using BioRender.com.

### 2.3 Oxycodone Self-Administration

Self-administration sessions were conducted in operant chambers (Med Associates, St. Albans, VT) equipped with two retractable levers, a stimulus cue light above each lever, and a sound-attenuating fan. Self-administration sessions began at the start of the dark phase of the dark/light cycle. The acquisition and escalation of oxycodone (Oxycodone HCl; B&B pharmaceutical, Englewood, CO) use was measured in a two-phased self-administration paradigm. The acquisition phase utilized a fixed-ratio 1 (FR1) reinforcement schedule in ten daily 2-h ShA sessions. A response at the active lever resulted in either an oxycodone infusion (0.15 mg/kg/infusion) or saline infusion (0.9%, 0.15 mL/infusion) delivered over 5-s and illumination of a cue light (7.5-W white light, 20-s). A 20-s timeout period (TO20) followed each infusion. Inactive lever responses were recorded but did not result in an infusion or cue light activation. The escalation phase consisted of a similar reinforcement schedule (FR1:TO20) to the acquisition phase. The escalation phase was conducted during ten daily 12-h LgA sessions with a 2-d break between sessions 5 and 6.

### 2.4 Behavioral Phenotypes

#### 2.4.1 Number of infusions and Total Intake

The number of infusions within each session was used to evaluate the overall pattern of intake during the acquisition and escalation phases. We used the number of infusions delivered within the first 15-m of each 2-h session to measure the initiation of oxycodone taking during each ShA session. The total number of infusions was also used to calculate total oxycodone intake (mg/kg) by multiplying the number of infusions by the unit dose (0.15 mg/kg/infusion) during each phase. These measures provided an overview of intake trends between the strains as previously reported (21).

#### 2.4.2 Lever Discrimination

Lever discrimination was used to determine the ability of animals to differentiate between the active lever and the inactive lever. This provides an assessment of learning during the early sessions of the acquisition phase. This measure was calculated by dividing the number of presses on the active lever that resulted in an infusion for a session by the total number of active and inactive lever presses. Inactive lever presses did not include responses during the 20-s timeout period following a press on the active lever.

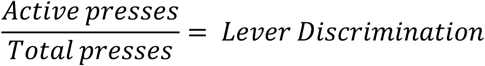

#### 2.4.3 Latency to the First Infusion

Latency to the first infusion was determined by recording the time the rat took to press the active lever following the onset of the session. This measure was used to characterize the initiation of each oxycodone self-administration session during the acquisition phase. The latency to initiate each session is anticipated to decrease as rodents learn the goal-directed behavior (32). Progressive decreases in latencies across sessions reflects the consolidation of the goal-directed behavior between sessions rather than within-session learning.

#### 2.4.4 Load Time

Drug self-administration sessions are often initiated with a drug loading phase (e.g., front-loading) in which rats engage in a high rate of responding at the beginning of self-administration sessions (33,34). The loading phase is an index of drug reinforcement and the regulation of the rat’s preferred drug concentrations (29,30). To quantify the drug loading phase, we calculated “Load Time” by subtracting the latency to the first infusion from the latency to deliver three infusions. This metric was utilized to characterize early session responding during the acquisition phase as a measure of drug regulation coinciding with learning.

#### 2.4.5 Escalation Metrics

An escalation index was utilized to determine the magnitude of escalation of oxycodone infusions throughout the escalation phase (24). The escalation index was calculated by averaging the total number of infusions during the last three LgA sessions and dividing by the average number of infusions from the first three sessions of LgA. Normalizing later intake to early intake controls for baseline differences between strains and allows for direct strain comparisons in the escalation of intake.

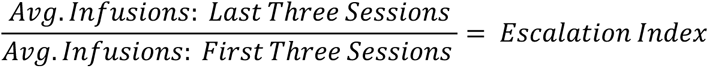

Regression-derived slopes of infusions across the escalation phase were used to complement the escalation index measure. The slope of infusions captures the rate of escalation and incorporates infusion data from all sessions rather than select “early” and “late” sessions but assumes that there is a linear relationship between session and number of infusions. A slope of 0 indicates stable intake while a positive slope indicates progressive escalation and a negative slope indicates reduced intake over time.

#### 2.4.6 Inter-infusion Interval

The inter-infusion interval was used as a measure of temporal organization of drug taking during the escalation phase (35). The average inter-infusion interval was calculated as the average time elapsed between successive infusions during the escalation phase. Progressive shortening of inter-infusion intervals across the escalation phase indicates that escalation of drug use coincides with within-session adaptations resulting from motivation, tolerance, and/or behavioral control (27,30,36).

#### 2.4.7 Burst Responding

Previous reports have indicated that rats engage in dysregulated drug intake by self-administering short bursts of infusions followed by a long drug-free interval (27,37–39). To capture “burst responding”, raw output data files containing the cumulative records of each timestamped behavioral event (e.g., active/inactive response, infusion, time-out responses) were used. The raw data from these files were extracted, and a custom MATLAB (Mathworks, R2025a, 25.1.0) script was used to determine the number of burst episodes, the average infusions per burst episode, inter-burst intervals, and the proportion of total infusions that occurred during a burst episode (40). A burst episode was defined as a minimum of 3 infusions occurring within a 90-s period, similar to previous reports (27,29). Once this minimum threshold was detected, the burst episode would continue until the rat no longer met the minimum criteria. This results in burst episodes of varying magnitude that we defined by the number of infusions per burst episode. We also determined the time elapsed between burst episodes (e.g., inter-burst intervals) analogous to inter-infusion intervals. Lastly, to determine how strains differentially distribute their infusions in burst episodes vs. single discrete events, we calculated the ratio of infusions within burst episodes to total infusions by dividing the number burst infusions by the total number of infusions for each rat. A higher ratio of burst responding would indicate that more infusions were self-administered as part of a burst episode.

### 2.5 Data Analysis

All data are presented as the mean ± standard error of the mean (SEM). Data were analyzed and graphed using both R (v.4.5.3) and GraphPad Prism (v.11.0.1). Prior to analysis, animals were excluded based on a set of predefined criteria. Rats were removed from the dataset if they experienced health complications (e.g., > 20% weight loss, infection, etc.), catheter failure, missing more than 3 sessions in either the acquisition or escalation phase, or failure to complete LgA session 10 of the escalation phase. If data from a single session was missing (e.g., computer failure, tether failure, animal health, etc.), that data point was imputed by calculating the mean of the immediately preceding and subsequent session. Summary measure data points that were greater than 2 standard deviations (SD) from the strain mean were considered outliers and winsorized as the strain mean ± 2 SD. One M520/N rat was excluded from all analyses because it failed to acquire the self-administration behavior (0 oxycodone infusions) during the acquisition phase.

Longitudinal measures collected across sessions (e.g., infusions, latency to 1^st^ infusion, burst episodes, etc.) were initially analyzed using comprehensive linear mixed-effects models. These included a global 4-way linear mixed-effects model with Strain, Group, and Sex as categorical factors, and Session as a continuous variable, as well as a reduced 3-way model excluding sex. Although these models provided an overall assessment of experimental effects, interpretation was limited by multiple higher-order interactions. Given this complexity, subsequent analyses focused on simplified models to better resolve biologically meaningful effects. Thus, separate 2-way linear mixed-effects models were conducted within each strain to examine: (1) the effect of Group (Saline v. Oxycodone) across Sessions, (2) the effect of Sex across Session, and (3) the interactive effects of Group and Session or Sex and Session. The Geisser–Greenhouse correction was used to correct for any potential deviations from sphericity. Significant interactions were followed by *post hoc* comparisons using a Tukey’s multiple comparison test to control for familywise error. To determine Strain differences, summary measures (e.g., total oxycodone intake, escalation index, average inter-infusion interval, regression-derived slopes, etc.) were analyzed using separate one-way ANOVAs within each Group (Saline or Oxycodone).

To further quantify changes in acquisition and escalation metrics (e.g., infusions, latency to 1^st^ infusion, load time, inter-infusion intervals, etc.), linear regression analyses were conducted for each subject with the slope of the regression line serving as an index of change. An F-test was used to determine whether the slope significantly differed from zero, and to compare slopes between Groups or Sex within each Strain. The escalation index was also analyzed using a one-sample t-test to determine if these measures differed from a theoretical value of 1 (no escalation). For all tests, statistical significance was set at p < 0.05.

## 3. Results

### 3.1 Acquisition: Infusions per Session

A self-administration paradigm (**Figure 1**) was utilized to characterize the acquisition of operant self-administration behavior reinforced with either oxycodone or saline in male and female rats from 7 inbred strains. Based on our prior work, we hypothesized that strains would differ in their oxycodone intake during the acquisition phase (21). Acquisition of self-administration behavior was characterized by a variety of measures during 10 daily ShA 2-h self-administration sessions. **Figure 2** illustrates the differences in oxycodone and saline infusions throughout the acquisition phase for each strain. Separate mixed-effect models were used for each strain to determine intake changed across the two groups. The M520/N, WKY/NCrl, and F344/NCrl strains displayed a significant interaction between Group and Session (M520/N: F_3.596, 100.7_ = 5.696, p < 0.001, WKY/NCrl: F_3.156, 91.52_ = 2.976, p = 0.033, F344/NCrl: F_2.291, 73.32_ = 12.73, p < 0.001). Further analysis of these interactions shows that Group differences emerged on later sessions in the acquisition phase suggesting greater oxycodone compared with saline intake (**Figure 2**). The LE/Stm strain was unique in that there was no difference between Groups and no significant interactions, but there was a significant main effect of Session (F_1.492, 35.80_ = 3.894, p = 0.041). This suggests that LE/Stm rats respond similarly regardless of oxycodone or saline reinforcement. In the LEW/Ss strain we observed a main effect of Group (F_1, 21_ = 13.51, p = 0.001), but no effect of Session or the interaction between Session and Group. This suggests that the LEW/Ss strain responds preferentially higher for oxycodone reinforcement, but there is little variation across the acquisition phase. Lastly, we observed no significant effects of Group, Session, or their interaction in the LEW/Crl and F344/Stm strains. This suggests that escalation of oxycodone or saline intake was not present in the LEW/Crl and F344/Stm strains during the acquisition phase. Linear regression analyses largely confirm these findings as indicated by significant positive slopes in the M520/N, WKY/NCrl, F344/NCrl, and LE/Stm strains acquiring oxycodone self-administration, while the saline self-administering animals displayed either non-significant slopes or a significantly negative slope in the LEW/Ss strain (**Table 2**).

**Figure 2:**
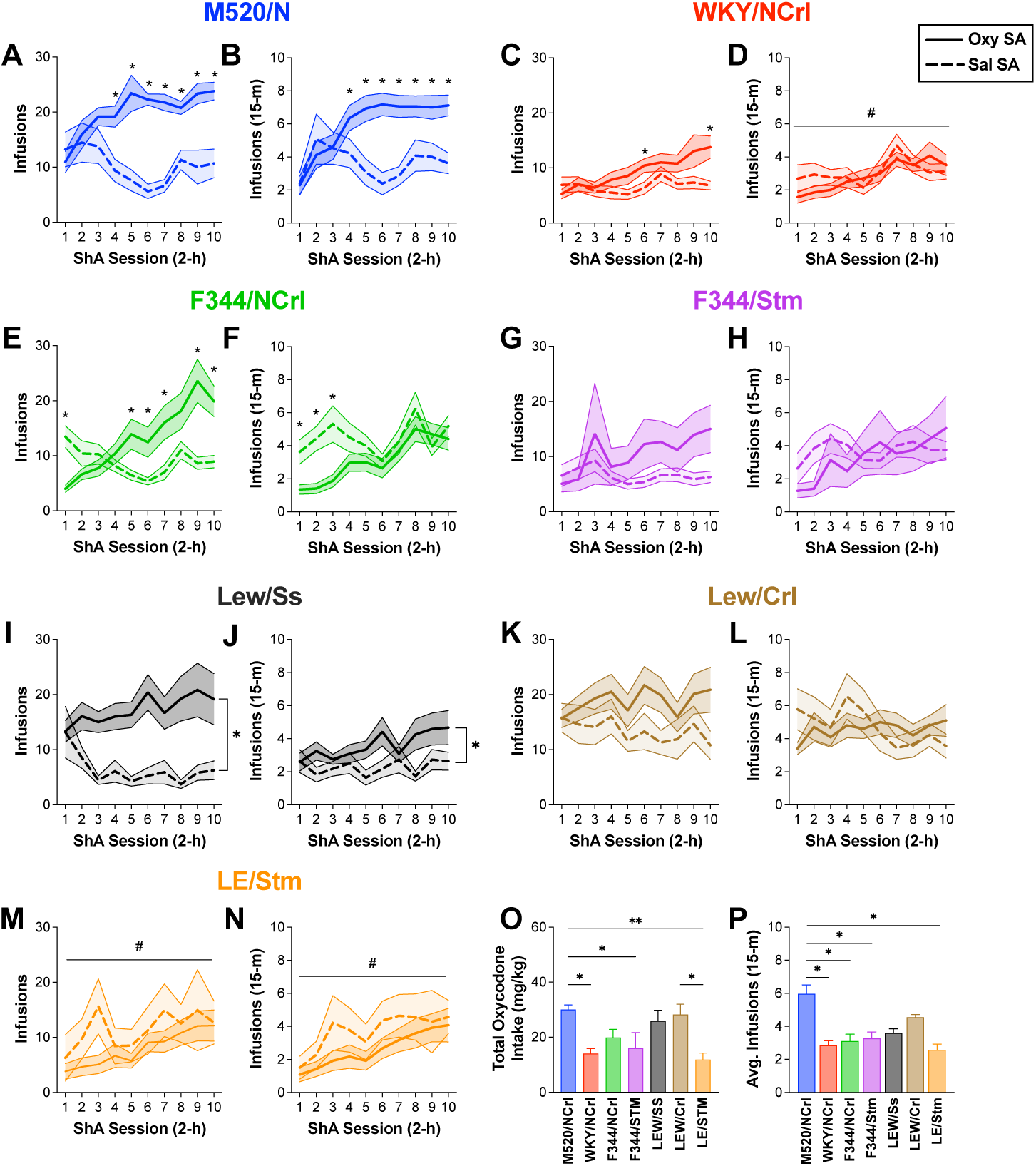
Self-administration in multiple inbred rat strains during the acquisition phase. Oxycodone (solid line) and saline (dashed line) infusions (mean ± SEM) during ten ShA (2-h) acquisition sessions are shown for the total session (**A, C, E, G, I, K, M**) and the first 15-m (**B, D, F, H, J, L, N**). A Tukey multiple comparison test detected several significant differences between Groups at individual Sessions in the M520/N (**A, B**), WKY/NCrl (**C**), and F344/NCrl (**E, F**) strains (* p < 0.05). WKY/NCrl rats also showed a significant increase in infusions across sessions regardless of Group (**D**, #, p < 0.05). Similarly, the LE/Stm strain displayed progressive increases in both total session (**M**) and first minutes (**N**) in both the saline and oxycodone groups (#, p <0.05). A main effect of Group was observed for the LEW/Ss strain in both total session (**I**) and first 15-m (**J**; *, p < 0.05), but neither indicated an interaction between Group and Session. When comparing the oxycodone treated group of each strain, we found that strains varied in both the total oxycodone intake (**O**) and the average intake during the first 15-m (**P**) with the M520/N rats showing significant differences compared with several other strains (*, p < 0.05; ** p < 0.01)

**Table 2.**
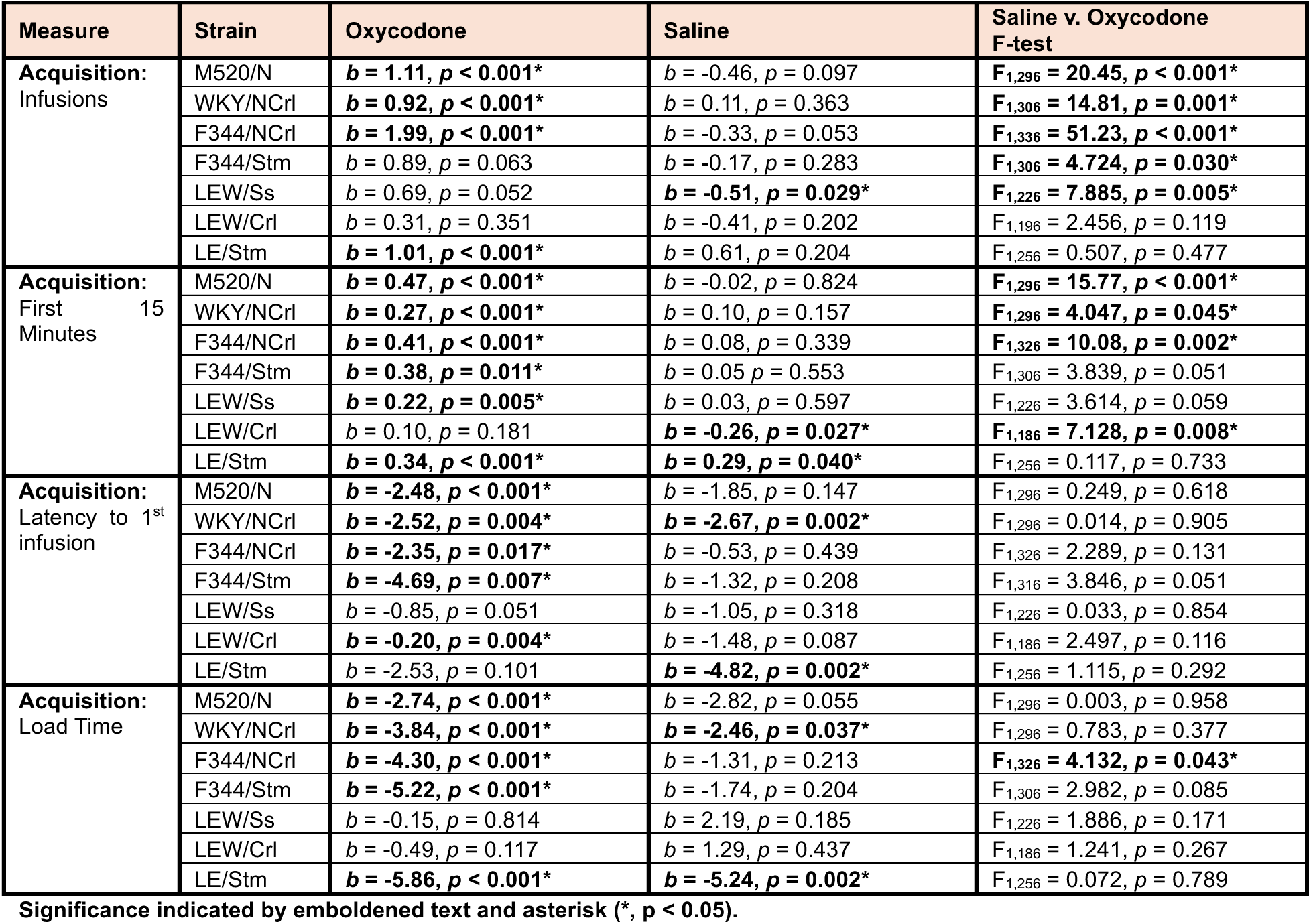
Linear regression analyses for longitudinal acquisition metrics.

To determine genetic differences in total oxycodone intake during the acquisition phase **(Figure 2O)**, a one-way ANOVA was conducted. As expected, oxycodone intake was significantly different between the strains (F_6, 90_ = 4.473, p < 0.001). Tukey post-hoc comparisons revealed that M520/N strain had significantly higher oxycodone intake compared to the WKY/NCrl, F344/Stm, and LE/Stm strains, while oxycodone intake in the LEW/Crl strain was higher than the LE/Stm strain (p < 0.05). Similar strain comparisons on total saline intake in the saline group revealed no significant differences (**Supplemental Figure 1A**).

### 3.2 Acquisition: Early Session Metrics

We also assessed how responding at the beginning of each acquisition session changed during the acquisition phase. We first analyzed the number of infusions during the first 15-m of each ShA session (**Figure 2**). Linear regression modeling indicated that all strains except the LEW/Ss strain significantly increased the number of oxycodone infusions administered in the first 15-m of each ShA session (**Table 2**). Every strain but one (LE/Stm) showed a decrease or no change in saline infusions in the first 15-min. A mixed-effect models revealed significant Group X Session interactions in the M520/N and F344/NCrl strains (M520/N: F_3.621, 91.32_ = 4.764, p = 0.003, F344/NCrl: F_2.704, 83.81_ = 3.211, p = 0.032). Further analysis of the interactions indicated significant differences between the oxycodone and saline groups in the early ShA sessions in the F344/NCrl strain while differences between the M520/N oxycodone and saline groups did not emerge until later in the acquisition phase. Among the other strains, a main effect of Group was observed for the LEW/Ss strain, with significantly higher overall oxycodone intake compared with saline during the first 15-m of ShA sessions (F_1, 21_ = 5.729, p = 0.026). A main effect of Session was observed in the WKY/NCrl and LE/Stm strains using a mixed-effect models, with both Groups in these strains increasing the number of infusions administered during the first 15-m (**Figure 2**; WKY/NCrl: F_4.539, 127.1_ = 3.959, p = 0.003, LE/Stm: F_1.651, 39.63_ = 4.968, p = 0.017). While mixed-effect models did not reveal any Group differences or interactions in the F344/Stm strain, the main effect of Session trended towards significance (p = 0.079). A one-way ANOVA was used to determine strain effects in the average number of infusions during the first 15-m **(Figure 2P)**. A main effect of Strain (F_6, 90_ = 3.889, p = 0.002) was observed with Tukey post hoc analyses revealing that the M520/N strain administered significantly more infusions during the first 15-m compared with the WKY/NCrl, F344/NCrl, F344/Stm, and LE/Stm strains. Similar strain comparisons on number of infusions during the first 15-m in the saline group revealed no significant differences (**Supplemental Figure 1B**).

The latency to the first infusion typically decreases across sessions reflecting consolidation of the goal-directed behavior between sessions. We hypothesized that the oxycodone Group would have shorter latencies compared to the saline treated groups, and that these latencies would progressively decrease during acquisition in a strain-dependent manner (**Figure 3**). Linear regression analyses support a progressive shortening in the latency to the first infusion in the oxycodone group of all strains except the LEW/Ss and LE/Stm strains (**Table 2**). Surprisingly, many of the strains in the saline group also displayed progressively shorter latencies with the WKY/NCrl and LE/Stm strains having significant, negative slopes on this measure (**Table 2**) and no significant differences in slopes between saline and oxycodone groups were observed in any strains. Mixed-effect models largely confirmed the regression modeling. No strain-specific models included a significant interaction effect between Session and Group. A main effect of Group in latency to the first infusion was only observed in the LEW/Ss strain (F_1, 21_ = 6.222, p = 0.021). The latencies to the first infusion decreased throughout acquisition in the M520/N, WKY/NCrl, F344/Stm, and LE/Stm strains as indicated by a significant main effect of Session (M520/N: F_2.039,57.10_ = 3.463, p = 0.037, WKY/NCrl: F_3.012, 84.34_ = 3.437, p = 0.020, F344/Stm: F_2.194, 65.81_ = 3.879, p = 0.022, LE/Stm: F_2.472, 59.34_ = 3.062, p = 0.044). The F344/NCrl, LEW/Crl, and LEW/Ss strains did not display significant differences between Groups or Sessions. Interestingly, the LEW/Crl and LEW/Ss strains initiate lever responding very quickly, even in the first few sessions (Figure 3). Despite the rapid initial responding, the LEW/Crl strain displayed a modest, but statistically significant negative slope, whereas the LEW/Ss strain maintained relatively stable responding throughout the acquisition phase (**Table 2**). A one-way ANOVA was used to analyze the average latency to first infusion throughout acquisition as a summary measure to assess genetic effects **(Figure 3O)**. We observed significant differences between strains in the average latency to the first infusion (F_6, 90_ = 4.544, p < 0.001) with the LEW/Crl, LEW/Ss, and M520/N showing the shortest latencies. Similar strain comparisons on the latency to first infusion in the saline group revealed a main effect of strain (F_6, 88_ = 2.987, p = 0.011). Tukey post hoc analysis confirms the LE/Stm strain had significantly longer latencies to the first saline infusion than the WKY/NCrl, F344/NCrl and LEW/Crl strains (**Supplemental Figure 1C**).

**Figure 3:**
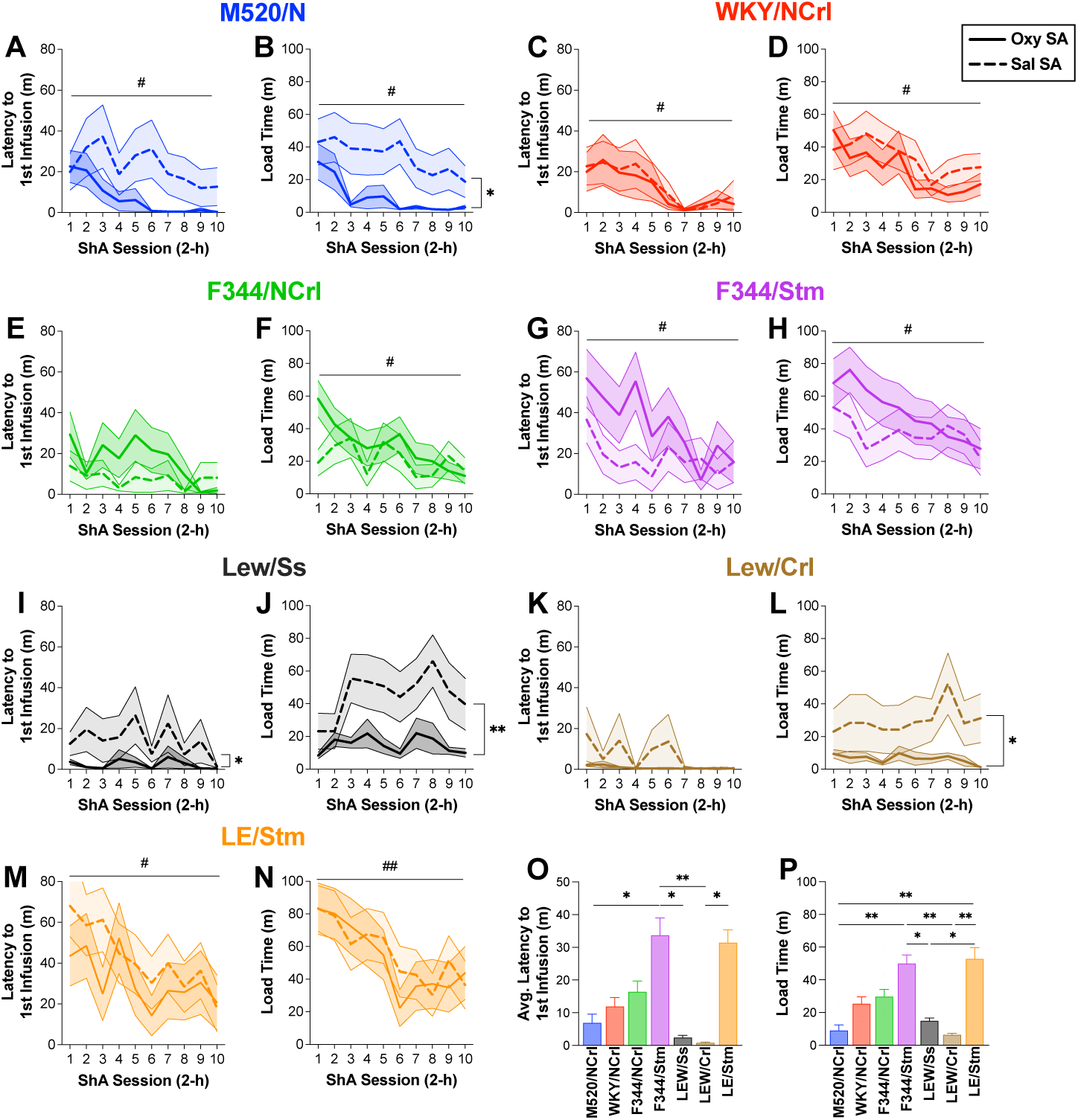
Early session measures of self-administration in multiple inbred rat strains during the acquisition phase. Latency to first infusion (**A, C, E, G, I, K, M**) and load time (**B, D, F, H, J, L, N**) are displayed for both the oxycodone (solid line) and saline (dashed line) Groups (mean ± SEM). Most strains displayed a progressive shortening of latency to first infusion, load time or both across the acquisition phase (# p < 0.05; #, p < 0.01). There were surprisingly few differences between the Groups except the LEW/Ss where the oxycodone group showed shorter latencies (**I**) and load times (**J**), and the M520/N (**B**) and LEW/Crl (**L**) that displayed shorter load times in the oxycodone group. When comparing oxycodone treated groups of each strain, there was significant strain variation in the latency to the first infusion (**O**) and load time (**P**) with the F344/Stm and the LE/Stm generally have slower initiation of oxycodone self-administration (* p < 0.05; ** p < 0.01; *** p < 0.001)

We characterized how rats regulated oxycodone intake at the beginning of the session by calculating the drug loading time. Drug loading is a phenomenon observed in IVSA studies in which rats engage in a high rate of responding at the beginning of self-administration sessions to rapidly reach desired drug levels (33,34). Linear regression modeling indicated that all, but the LEW/Crl and LEW/Ss, strains showed significant shortening of oxycodone loading times throughout the acquisition phase while only the LE/Stm and WKY/NCrl strains also showed significant shorting of saline loading times (**Table 2**). The difference in slopes between the oxycodone group and the saline group was only significant for the F344/NCrl strain. Using mixed-effect models, no strains indicated a significant interaction between Group and Session, but several strains displayed a main effect of the Group with the oxycodone group displaying significantly shorter load times compared to the saline group (M520/N: F_1, 28_ = 14.33, p = 0.018, LEW/Ss: F_1, 21_ = 11.07, p = 0.003, LEW/Crl: F_1, 17_ = 9.261, p = 0.007). No significant differences between Groups were found in the WKY/NCrl, F344/Stm, F344/NCrl, and LE/Stm strains. A main effect of Session was observed in the M520/N, F344/NCrl, and LE/Stm strains, indicating that load times significantly decreased throughout acquisition regardless of Group (M520/N: F_2.923, 81.84_ = 3.599, p = 0.018, F344/NCrl: F_2.224, 68.94_ = 5.063, p = 0.007, LE/Stm: F_1.779, 42.70_ = 6.419, p = 0.005). The average load time during oxycodone acquisition was analyzed using a one-way ANOVA to identify genetic differences between the strains **(Figure 3P)**. Results revealed a significant strain difference with the LEW/Crl, LEW/Ss, and M520/N strains showing the shortest load times (F_6, 90_ = 5.589, p < 0.001). Similar strain comparisons on the load time in the saline group revealed no significant differences (**Supplemental Figure 1D**).

### 3.3 Acquisition: Lever Discrimination

To further characterize how the strains acquired lever responding with oxycodone or saline reinforcement, we investigated the ability of each strain to discriminate between the active (oxycodone/saline-paired) and the inactive lever (**Supplementary Figure 2**). Using a lever discrimination index (% Responding for Active Lever), mixed-effect models were used to determine differences between Groups and Sessions during acquisition for each strain. Although no strains indicated a significant interaction between Group and Session, we found a main effect of Group in the WKY/NCrl and LEW/Crl, with the oxycodone group showing greater active lever discrimination compared with the saline group (WKY/NCrl: F_1, 29_ = 7.288, p = 0.012, LEW/Crl: F_1, 17_ = 7.535, p = 0.014). No Group differences were found in the lever discrimination for any of the other strains. Active lever discrimination improved in the WKY/NCrl, F344/NCrl, F344/Stm, and LE/Stm strains as indicated by a main effect of Session (WKY/NCrl: F_4.361, 126.5_ = 4.083, p = 0.003, F344/NCrl: F_3.011, 93.34_ = 4.016, p = 0.010, F344/Stm: F_4.073, 118.1_ = 3.317, p = 0.013, LE/Stm: F_3.120, 74.87_ = 2.730, p = 0.048). No significant session differences were found in the other strains.

### 3.4 Escalation: Infusions per Session

To facilitate escalation of oxycodone intake, all rats underwent 10 daily LgA (12-h) self-administration sessions. Linear regression modeling confirms that all, but the F344/Stm strain, displayed a significant escalation in oxycodone intake while saline intake significantly decreased during the escalation phase in all strains, but the LEW/Crl (**Table 3**; **Figure 4**). Mixed-effect models conducted for each strain confirmed these results with a significant interaction between Group and Session for the M520/N, WKY/NCrl, F344/NCrl, LEW/Ss, LEW/Crl, and LE/Stm strains (M520/N: F_3.115, 87.23_ = 12.02, p < 0.001, WKY/NCrl: F_1.471, 41.18_ = 6.793, p = 0.006, F344/NCrl: F_2.198, 68.14_ = 10.11, p < 0.001, LEW/Ss: F_1.738, 36.49_ = 4.584, p = 0.021, LEW/Crl: F_2.656, 45.16_ = 6.073, p = 0.002, LE/Stm: F_1.857, 44.58_ = 16.25, p < 0.001). Further analysis confirmed that differences between Groups emerged early in the escalation phase in the M520/N, WKY/NCrl, F344/NCrl, LEW/Ss, and LEW/Crl strains while differences occurred in the second half of the escalation phase in the LE/Stm strain (**Figure 4**). The F344/Stm strain displayed a main effect of Group, with consistently higher active lever responding in the oxycodone group throughout the escalation phase (F344/Stm: F_1, 29_ = 23.81, p < 0.001). Interestingly, the F344/Stm strain did not escalate oxycodone intake as demonstrated by a non-significant slope estimate in linear regression modeling and no significant interaction effect or Session effect in the mixed effects ANOVA. We also assessed the total amount of oxycodone intake during the escalation phase as a summary measure **(Figure 4H).** A one-way ANOVA in rats from the oxycodone treatment group found significant differences between strains (F_6, 90_ = 7.532, p < 0.001). Tukey post hoc analysis revealed the M520/N strain had significantly higher intake than all other strains (p < 0.05), suggesting that the M520/N is uniquely susceptible to high oxycodone intake compared with the other strains tested. Similar strain comparisons in total infusions for the saline group revealed strain differences, with Tukey post hoc analysis finding the M520/N administered significantly more saline than the F344/Stm and LEW/Ss strains (**Supplemental Figure 3A**).

**Figure 4:**
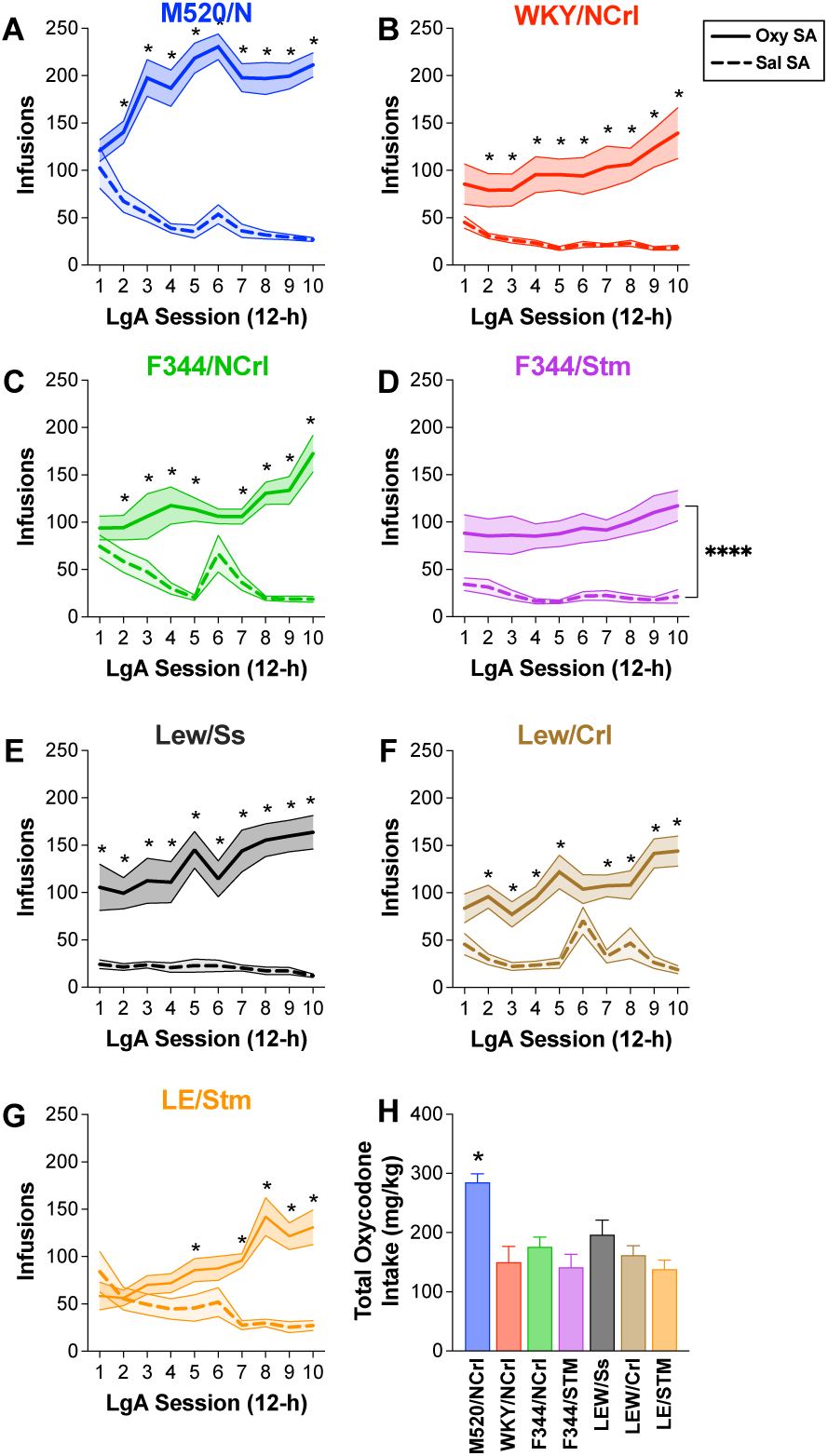
Self-administration in multiple inbred rat strains during the acquisition phase. Total infusions (mean ± SEM) across ten 12-h escalation sessions are shown for oxycodone (solid line) and saline (dashed line) Groups in the M520/N (**A**), WKY/NCrl (**B**), F344/NCrl (**C**), F344/Stm (**D**), LEW/Ss (**E**), LEW/Crl (**F**), and LE/Stm (**G**) strains. Tukey multiple comparison test detected several significant differences between Groups at individual Sessions in all strains except F344/Stm, as indicated by a filled circle at that data point (p < 0.05). Total infusions were significantly higher in the oxycodone group compared with the saline group in the F344/Stm strain (**** p < 0.0001). When comparing the oxycodone group of each strain, we found total oxycodone intake (**H**) was highest in the M520/N strain compared with all other strains (* p < 0.05).

**Table 3.** Linear regression analyses for longitudinal escalation metrics.

| Measure | Strain | Oxycodone | Saline | Saline v. Oxycodone F-test |
| --- | --- | --- | --- | --- |
| **Escalation: Infusions** | M520/N | **$b = 7.69, p < 0.001^*$** | **$b = -6.34, p < 0.001^*$** | **$F_{1,296} = 39.95, p < 0.001^*$** |
|  | WKY/NCrI | **$b = 5.78, p = 0.008^*$** | **$b = -2.12, p < 0.001^*$** | **$F_{1,296} = 15.00, p < 0.001^*$** |
|  | F344/NCrI | **$b = 6.45, p < 0.001^*$** | **$b = -5.17, p < 0.001^*$** | **$F_{1,326} = 33.10, p < 0.001^*$** |
| | F344/Stm | $b = 3.19, p = 0.066$ | **$b = -1.24, p = 0.035^*$** | **$F_{1,306} = 6.527, p = 0.013^*$** |
|  | LEW/Ss | **$b = 7.44, p < 0.001^*$** | **$b = -1.03, p = 0.033^*$** | **$F_{1,226} = 13.58, p < 0.001^*$** |
| | LEW/CrI | **$b = 6.30, p < 0.001^*$** | $b = -0.43, p = 0.695$ | **$F_{1,186} = 11.81, p < 0.001^*$** |
|  | LE/Stm | **$b = 9.35, p < 0.001^*$** | **$b = -5.24, p < 0.001^*$** | **$F_{1,256} = 57.12, p < 0.001^*$** |
| **Escalation: Inter-infusion intervals** | M520/N | **$b = -0.27, p < 0.001^*$** | **$b = 1.96, p < 0.001^*$** | **$F_{1,296} = 45.73, p < 0.001^*$** |
|  | WKY/NCrI | **$b = -0.79, p < 0.001^*$** | **$b = 2.65, p < 0.001^*$** | **$F_{1,303} = 37.08, p < 0.001^*$** |
|  | F344/NCrI | **$b = -0.57, p < 0.001^*$** | **$b = 4.25, p < 0.001^*$** | **$F_{1,326} = 43.39, p < 0.001^*$** |
| | F344/Stm | $b = -0.07, p = 0.937$ | **$b = 4.12, p = 0.006^*$** | **$F_{1,301} = 5.231, p = 0.023^*$** |
|  | LEW/Ss | **$b = -0.66, p < 0.001^*$** | **$b = 2.99, p = 0.003^*$** | **$F_{1,226} = 14.32, p = 0.002^*$** |
| | LEW/CrI | **$b = -0.61, p < 0.001^*$** | $b = 1.48, p = 0.295$ | $F_{1,186} = 2.432, p = 0.121$ |
|  | LE/Stm | **$b = -0.79, p = 0.001^*$** | **$b = 2.46, p < 0.001^*$** | **$F_{1,252} = 22.95, p < 0.001^*$** |
| **Escalation: Burst Episodes** | M520/N | **$b = 1.35, p < 0.001^*$** | **$b = -0.83, p < 0.001^*$** | **$F_{1,296} = 44.82, p < 0.001^*$** |
| | WKY/NCrI | $b = 0.36, p = 0.098$ | **$b = -0.11, p < 0.001^*$** | **$F_{1,296} = 5.258, p = 0.023^*$** |
|  | F344/NCrI | **$b = 0.70, p = 0.003^*$** | **$b = -0.67, p < 0.001^*$** | **$F_{1,326} = 21.98, p < 0.001^*$** |
| | F344/Stm | $b = 0.25, p = 0.094$ | $b = -0.10, p = 0.149$ | **$F_{1,306} = 4.726, p = 0.031^*$** |
| | LEW/Ss | $b = 0.59, p = 0.059$ | $b = -0.04, p = 0.432$ | $F_{1,226} = 3.725, p = 0.055$ |
| | LEW/CrI | $b = 0.30, p = 0.343$ | $b = 0.06, p = 0.692$ | $F_{1,186} = 0.436, p = 0.510$ |
|  | LE/Stm | **$b = 0.58, p = 0.007^*$** | **$b = -0.64, p < 0.001^*$** | **$F_{1,256} = 22.49, p < 0.001^*$** |

Significance indicated by emboldened text and asterisk ( $* p < 0.05$ ).

### 3.5 Escalation: Escalation Metrics

To characterize escalation of oxycodone intake (oxycodone treatment group only), we utilized two complementary, yet distinct, metrics calculated based on self-infusions during the escalation phase **(Figure 5)**. The escalation index was used to measure the magnitude of change from the early escalation phase (Sessions 1-3) compared with the late escalation phase (Sessions 8-10). Surprisingly, no significant differences were found in the escalation index between the Strains despite significant differences in total oxycodone intake (F_6, 89_ = 0.7606, p = 0.603). This suggests that when controlling for baseline differences between strains, all strains increase their oxycodone intake similarly during the escalation phase. Intake escalation was confirmed using a one-sample t-test to determine if the average escalation index for each strain was significantly different than a hypothetical magnitude of one (i.e., no change/escalation). All strains displayed significant intake escalation (**Figure 5A**). Using the slopes of self-infusions calculated in linear regression modeling **(Figure 5B and Table 3)**, we did not detect significant Strain differences in the rate of escalation (F_6, 91_ = 0.9781, p = 0.445), suggesting similar intake escalation across with positive slopes for all strains except the F344/Stm strain. Using similar metrics to evaluate changes in saline intake revealed significant reduction in saline intake in most strains and negative slopes of saline intake that differed in a strain dependent manner (F_6, 87_ = 4.625, p < 0.001). Specifically, saline treated M520/N rats had a larger negative slope than the F344/NCrl, LEW/Crl, and LEW/Ss strains based on Tukey post-hoc analysis (**Table 3; Supplemental Figure 3B-C**).

**Figure 5:**
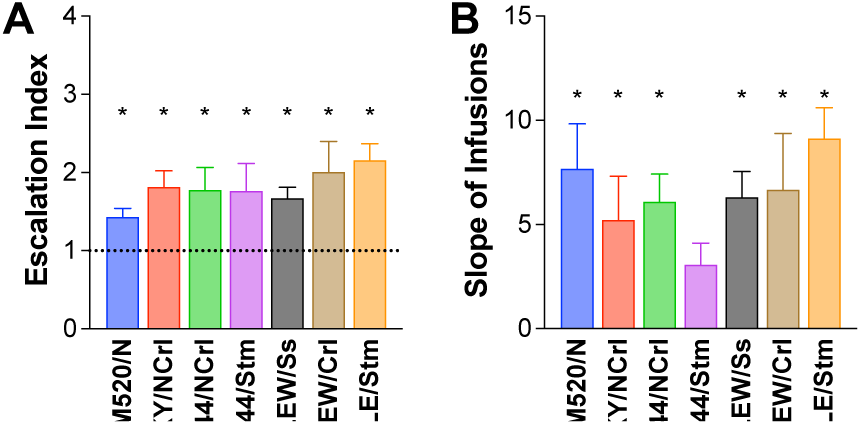
Change in intake during the escalation phase in multiple inbred rat strains. (**A**) Escalation index for the oxycodone group in each strain. No strain differences were detected (p = 0.6028). However, all strains showed escalation indices that were significantly greater than the hypothetical mean of 1 using a one-sample t-test (* p < 0.05). (**B**) The slope of infusions across LgA sessions was used to measure the rate of change during escalation. Slopes did not significantly differ between the strains (p = 0.4447), but linear regression modeling indicated that all strains except F344/Stm showed significance (* p < 0.05; **Table 3**).

### 3.5 Escalation: Inter-Infusion Interval

To capture the within-session temporal organization of self-administration behavior, we assessed the average inter-infusion intervals for each session during the escalation phase. Progressive shortening of inter-infusion intervals across the escalation phase indicates that escalation of drug use coincides with within-session adaptations in the regulation of drug intake. Linear regression modeling supports a progressive shortening of inter-infusion intervals across sessions for all strains except the F344/Stm in the oxycodone group as indicated by significant negative slopes (**Table 3**). Rats self-administering saline in all strains displayed significant positive slopes indicated progressive lengthening of inter-infusion intervals, with the exception of the LEW/Crl (**Table 3**). Using a mixed-effect model for each strain, a significant difference between Groups was found for all strains, with the oxycodone group having shorter inter-infusion intervals (**Figure 6**). The M520/N, WKY/NCrl, F344/NCrl, and LE/Stm strains displayed a significant interaction between Group and Sessions, with the oxycodone group showing progressively shorter inter-infusion intervals compared with the saline group that had stable or increasing inter-infusion intervals over the course of the escalation phase (M520/N: F_1.981, 55.47_ = 13.45, p < 0.001, WKY/NCrl: F_2.124, 60.88_ = 11.53, p < 0.001, F344/NCrl: F_3.077, 95.38_ = 9.438, p < 0.001, LE/Stm: F_2.602, 61.28_ = 5.527, p = 0.003). A main effect of Group was found in the F344/Stm, LEW/Ss, and LEW/Crl strains, with the oxycodone group in these strains engaging in significantly shorter inter-infusion intervals (F344/Stm: F_1, 29_ = 16.91, p < 0.001, LEW/Ss: F_1, 21_ = 58.41, p < 0.001, LEW/Crl: F_1, 17_ = 11.83, p = 0.003). A one-way ANOVA was used to determine strain effects in the average inter-infusion interval for the oxycodone group **(Figure 6H)**. A main effect of Strain was found in the average inter-infusion interval during the escalation phase (F_6, 90_ = 3.222, p = 0.007). Tukey post hoc analysis revealed that the F344/Stm strain had a significantly higher average inter-infusion interval compared with the M520/N and F344/NCrl strains (p < 0.05). Similar strain comparisons in the saline group revealed significant differences in the inter-infusion interval between strains (F_6, 87_ = 4.407, p < 0.001) (**Supplemental Figure 3D**). Specifically, saline treated F344/Stm rats had a significantly higher inter-infusion interval compared to the M520/N and LE/Stm strains based on Tukey post-hoc analysis.

**Figure 6:**
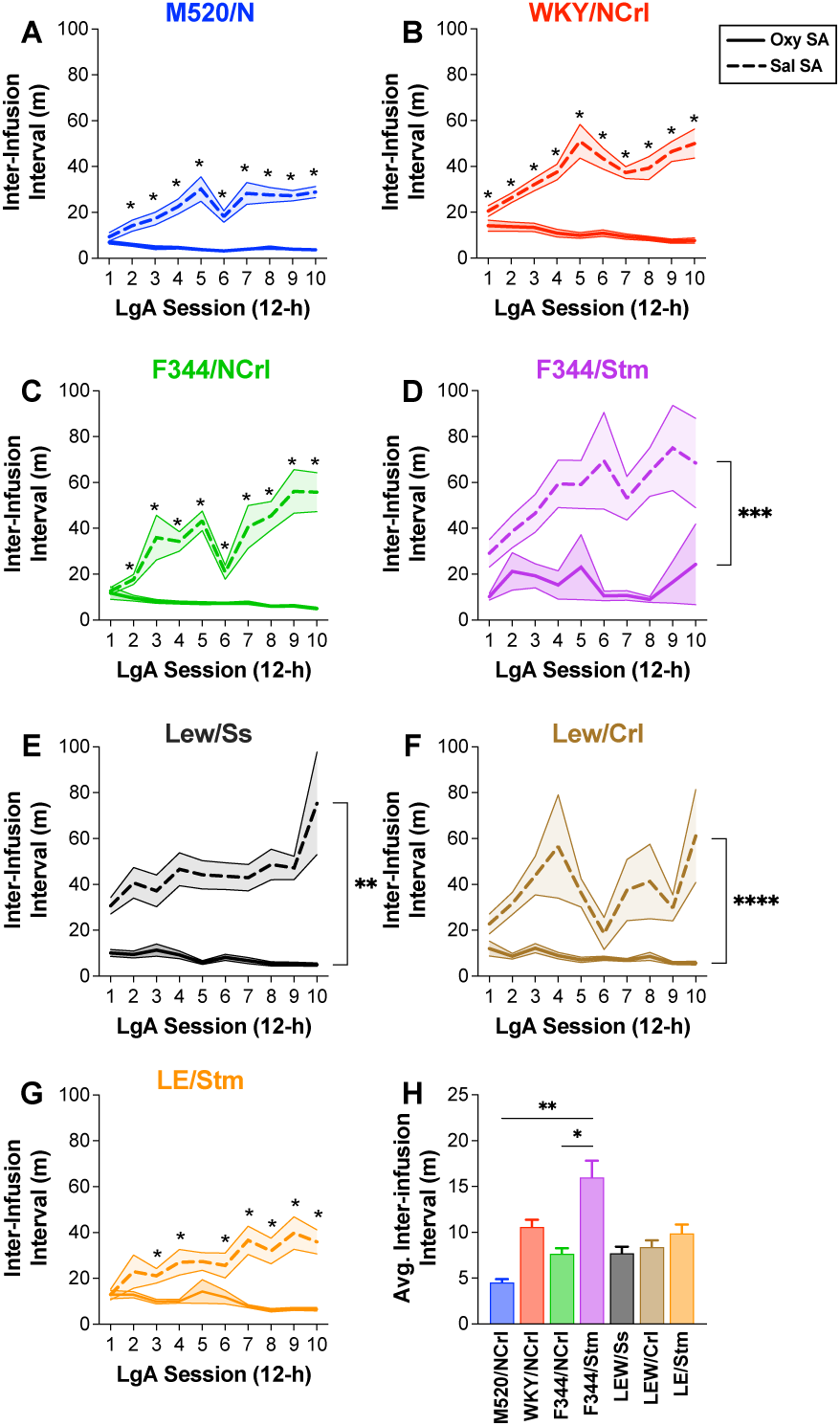
Temporal organization of infusions during the escalation phase in multiple inbred rat strains. Inter-infusion intervals (mean ± SEM) across escalation sessions are shown for oxycodone (solid line) and saline (dashed line) Groups in the M520/N (**A**), WKY/NCrl (**B**), F344/NCrl (**C**), F344/Stm (**D**), LEW/Ss (**E**), LEW/Crl (**F**), and LE/Stm (**G**) strains. Tukey multiple comparison test detected several significant differences between Groups at individual Sessions in the M520/N, WKY/NCrl, F344/NCrl, and LE/Stm strains (* p < 0.05). The F344/Stm, LEW/Ss, and LEW/Crl strains showed significant main effects of Group (** p < 0.01; *** p < 0.001; **** P < 0.0001). (**H**) The average inter-infusion interval during escalation revealed a main effect of strain with F344/Stm displaying longer intervals compared with M520/N and F344/NCrl (*, p < 0.05; ** p < 0.01).

### 3.6 Escalation: Burst Episodes

Based on our assessment of within-session inter-infusion intervals, we were interested in additional metrics that may reflect dysregulated oxycodone intake and vary between strains within the escalation phase. Previous reports indicated that rats engage in dysregulated drug intake by self-administering short bursts of infusions followed by a long drug-free interval (27,37–39). We observed similar patterns in some strains but not others throughout the escalation phase as depicted in the raster plots of representative rats from each strain (**Fig. 7**).

**Figure 7:**
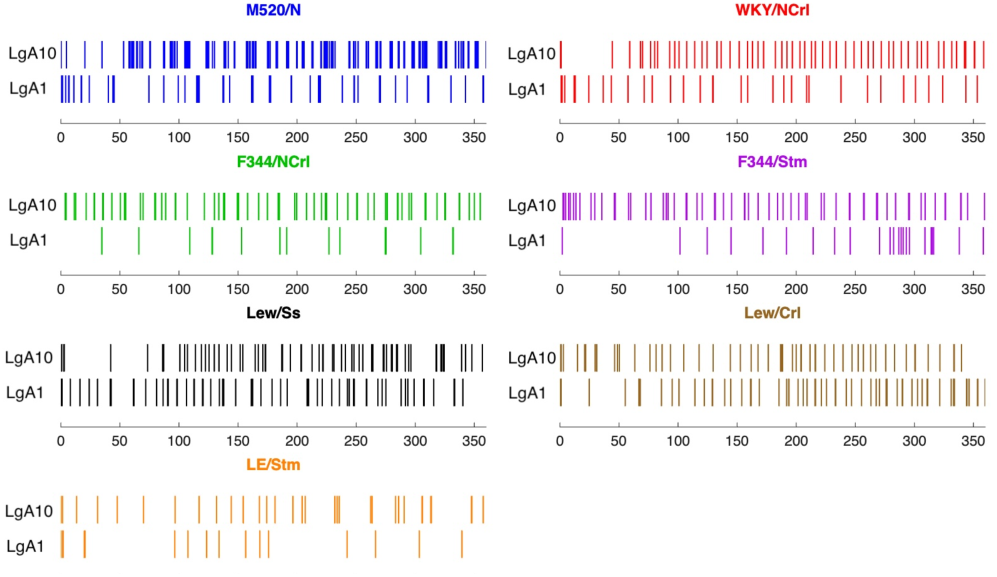
Representative within-session patterns of oxycodone intake during the escalation phase in multiple inbred rat strains. Raster plots show the timing of individual oxycodone infusions from representative animals of each strain during the 1st and 10th LgA sessions of the escalation phase.

Linear regression modeling indicates that only a few strains in the oxycodone group (i.e., M520/N, F344/NCrl, LE/Stm) show a significant, progressive increase in burst responding, while most of the strains in the saline group show either no change or a significant decrease in burst responding (**Table 3**). A mixed-effect model detected similar effects with a significant interaction between Group and Session in the M520/N, F344/NCrl, LEW/Crl, and LE/Stm strains (M520/N: F_3.722, 104.2_ = 10.80, p < 0.001, F344/NCrl: F_1.980, 61.38_ = 7.424, p = 0.001, LEW/Crl: F_2.148, 36.52_ = 3.290, p = 0.045, LE/Stm: F_2.631, 63.14_ = 5.314, p = 0.004). The oxycodone groups in the F344/NCrl and LE/Stm strains had a significantly higher number of burst episodes in the later escalation sessions while group differences in the M520/N and LEW/Crl strains emerged early in the escalation phase (**Figure 8**). A main effect of Group in the WKY/NCrl, F344/Stm, and LEW/Ss strains indicates a significantly higher number of burst episodes in the oxycodone group throughout the escalation phase (WKY/NCrl: F_1, 28_ = 10.62, p = 0.003, F344/Stm: F_1, 29_ = 10.25, p = 0.003, LEW/Ss: F_1, 21_ = 12.72, p = 0.002). Moreover, we observed very few burst episodes in the saline group suggesting that burst responding in our paradigm is a pattern of self-administration unique to oxycodone. Using a one-way ANOVA, we detected significant strain differences in the average number of burst episodes per session during the escalation phase (**Figure 8H**, F_6, 90_ = 11.76, p < 0.001) among the oxycodone consuming rats. Notably, the M520/N strain had a significantly higher number of burst episodes compared with all other strains suggesting that the M520/N strain is uniquely vulnerable to developing this dysregulated pattern of oxycodone intake. Despite burst responding being uncommon and inconsistent in the saline group, strains displayed significant differences in burst responding for saline (F_6, 63_ = 4.10, p = 0.002). The LE/Stm and F344/NCrl strains showed the highest average number of burst episodes (**Supplementary Figure 4A**).

**Figure 8:**
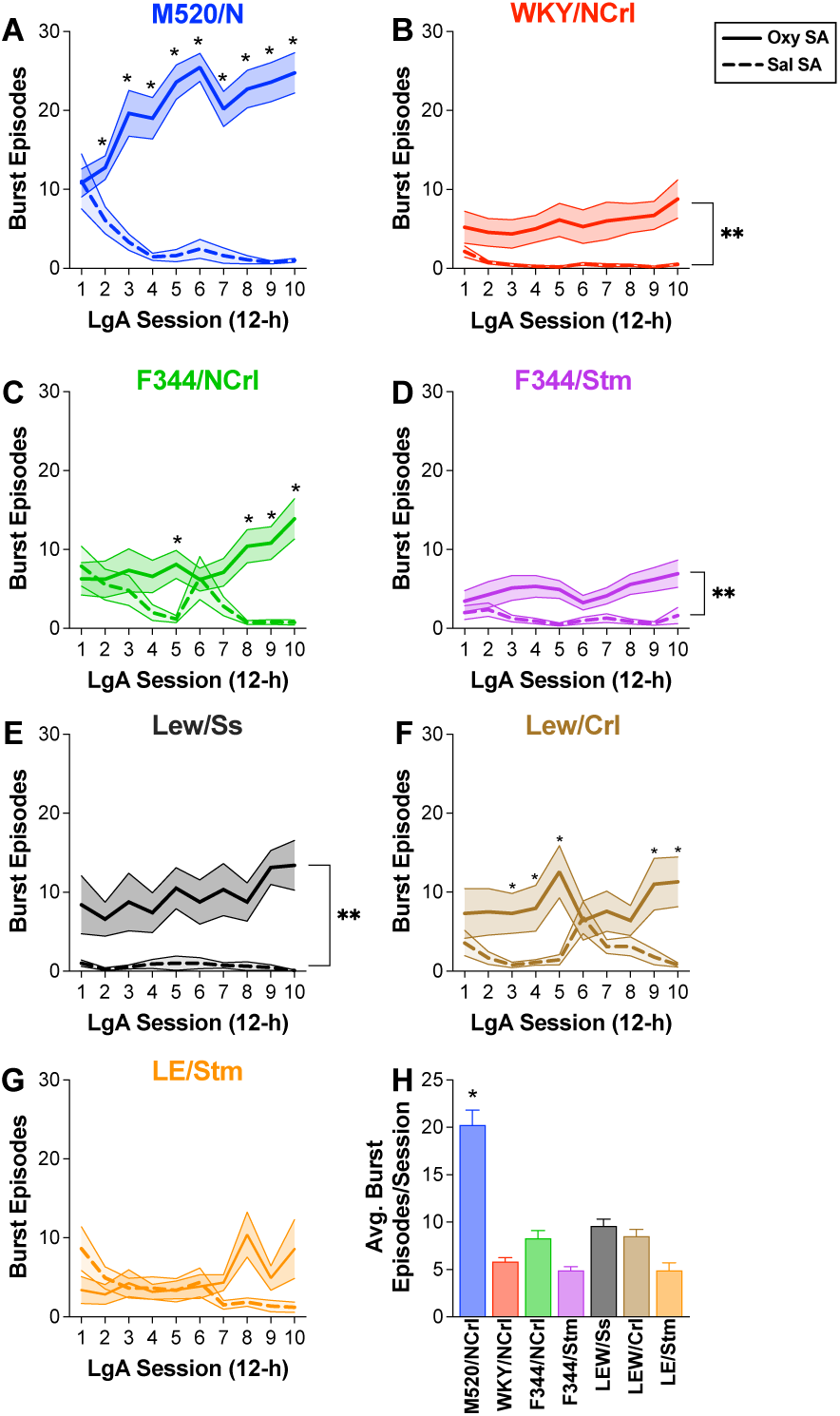
Burst responding during the escalation phase in multiple inbred rat strains. The number of burst episodes (mean ± SEM) per LgA session during the escalation phase is displayed for oxycodone (solid line) and saline (dashed line) Groups in the M520/N (**A**), WKY/NCrl (**B**), F344/NCrl (**C**), F344/Stm (**D**), LEW/Ss (**E**), LEW/Crl (**F**), and LE/Stm (**G**) strains. Tukey multiple comparison test detected several significant differences between Groups at individual Sessions in the M520/N, F344/NCrl, LEW/Crl, and LE/Stm strains, as indicated by a filled circle at that data point (p < 0.05). WKY/NCrl, F344/Stm, and LEW/Ss strains showed a significant main effect of Group (** p < 0.01). (**H**) Average number of burst episodes per session during escalation revealed a main effect of with the M520/N strain displaying significantly more burst episodes compared with all other strains (* p < 0.0005).

### 3.7 Escalation: Burst Measures

We were interested in identifying the proportion of oxycodone self-infusions that occurred within a burst episode during the escalation phase. We quantified this by calculating a Burst Ratio (% Burst Infusions). Strains differed significantly in the proportion of infusions delivered within a burst episode with the M520/N strain showing considerably higher ratio of burst infusions compared with all other tested strains except for the F344/NCrl strain (**Figure 9A**; F_6, 90_ = 4.492, p < 0.001). Strain differences in the proportion of saline infusions occurring during a burst episode were also observed, with the LE/Stm rats displaying higher proportion of saline infusions occurring during a burst episode compared to the WKY/NCrl and LEW/Ss strains (**Supplementary Figure 4B;** F_6, 88_ = 3.86, p = 0.002). To further characterize burst responding during escalation, we calculated the average magnitude of each burst by determining the number of infusions within each burst during each session. Strains did not differ in the number of infusions per burst (**Figure 9B**; F_6, 90_ = 1.397, p = 0.225). We also calculated the inter-burst interval to identify whether strains differed in the temporal organization of burst episodes throughout each session. There was a significant effect of Strain with the M520/N strain displaying the shortest inter-burst intervals (**Figure 9C**; F_6, 90_ = 3.085, p = 0.009), confirming more frequent burst episodes. In contrast, strains such as the F344/Stm exhibited much longer inter-burst intervals (∼200 min). Together, these findings indicate that oxycodone intake in the M520/N strain occurs in more frequent bursts of infusions, reflecting a distinct microstructural pattern of intake that may indicate dysregulated and/or compulsive drug intake.

**Figure 9:**
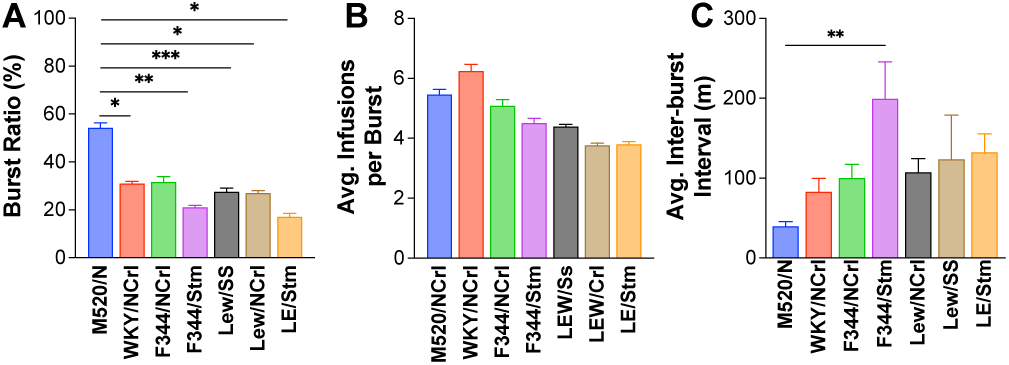
Characterization of burst responding for oxycodone during the escalation phase in multiple inbred rat strains. (**A**) The average ratio of infusions occurring in bursts was significantly higher in M520/N rats compared with all strains except F344/NCrl (* p < 0.05; ** p < 0.01; *** p < 0.001). and total bursts is displayed. (**B**) The average number of infusions per burst episode varied between strains but was not significantly different. (**C**) The average inter-burst interval for each strain varied between strains with the M520/N strain displaying the shortest average inter-burst interval (** p < 0.01).

### 3.8 Sex Differences

Mixed-effect models were utilized to determine if sex differences were present in several acquisition measures across the ShA sessions including total infusions, infusions in the first 15-m of the session, latency to the 1^st^ infusion, and load time **(Figure 10)**. Here, only rats in the oxycodone group were analyzed and we only report significant main effects of Sex or the interaction between Sex and Session. No significant main effect of Sex or Sex-Session interaction was observed with the number of infusions during acquisition. WKY/NCrl males displayed a statistical trend towards more infusions than WKY/NCrl females across the acquisition phase (F_1,12_ = 3.48, p = 0.087). Regression modeling indicated that both males and females from nearly all strains demonstrated significant linear increases in oxycodone infusions during the acquisition period (**Table 4**). Statistical comparison of the slopes between male and female WKY/NCrl rats indicates a significantly steeper slope in the male WKY/NCrl rats (**Table 4**). Analysis of oxycodone infusions within the first 15-m detected no significant main effects of Sex or Sex-Session interactions. However, linear regression modeling indicates significantly steeper slopes in female M520/N and F344/NCrl rats compared with male counterparts (**Table 4**). In contrast, male LEW/Ss and LE/Stm rats displayed significantly steeper slopes compared to females (**Table 4**). Analysis of the latency to the 1^st^ infusion revealed that male M520/N rats displayed significantly faster latencies to the 1^st^ infusion, particularly during the first two sessions, as demonstrated by a significant Sex-Session interaction (F_9, 135_ = 2.59, p = 0.009). A significant interaction between Sex and Session was also observed in the LEW/Ss strain, with males having a significantly longer latency to the 1^st^ infusion only during the first session (F_9, 90_ = 2.740, p = 0.007). Linear regression analyses confirmed these sex differences with female M520/N having steeper negative slopes compared with males, and male LEW/Ss rats having steeper negative slopes than females (**Table 4**). Analysis of load time revealed that female M520/N and F344/NCrl rats displayed significantly longer load times compared with males, as demonstrated by a significant Sex X Session interaction (M520/N: F_9,135_ = 3.74, p < 0.001; F344/NCrl: F_9,62_ = 2.24, p = 0.031). Linear regression modeling confirmed these results as female M520/N and F344/NCrl rats displayed steep negative slopes compared with males (**Table 4**). Together, these results indicate subtle differences between male and female rats in the early session latency measures where M520/N and WKY/NCrl males appear to initiate responding for oxycodone reinforcement more rapidly than females.

**Figure 10:**
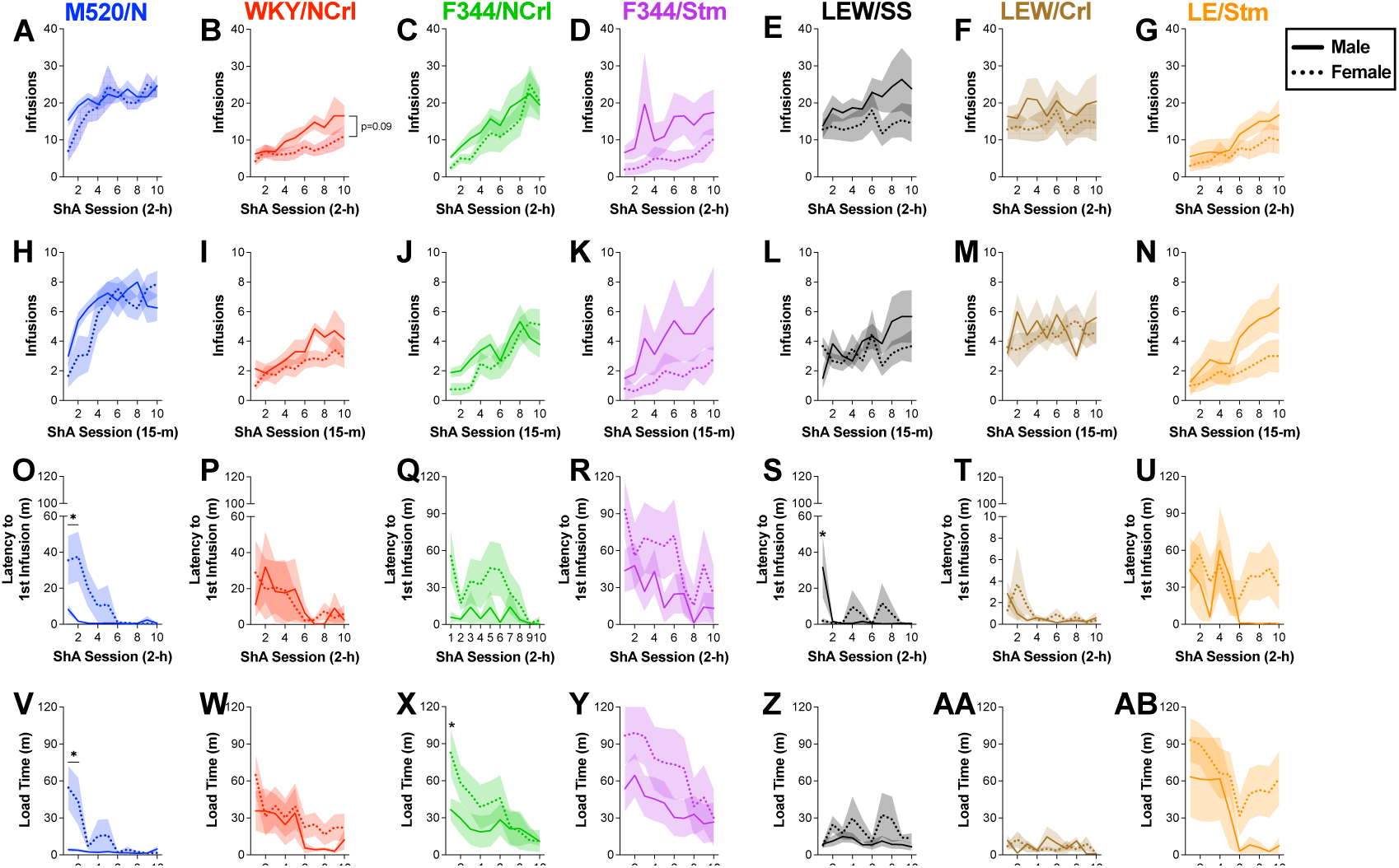
Comparison between male and females in acquisition metrics. Total infusions per 2-h session during the acquisition phase is displayed for males (solid line) and females (dashed line) in the M520/N **(A)**, WKY/NCrl **(B)**, F344/NCrl **(C)**, F344/Stm **(D),** LEW/Ss **(E)**, LEW/Crl **(F)**, and LE/Stm **(G)** strains. No significant differences in infusions between the sexes were observed in any of the strains, although WKY/NCrl males showed a trend toward higher intake than females (p = 0.09). No differences between males and females were observed in the number of infusions during the first 15-m of a session is displayed for males (solid line) and females (dashed line) in all strains (panels **H-N**). The average latency to the first infusion is shown in panels **O-U**. Tukey multiple comparison test detected several subtle differences between males and females at individual Sessions in the M520/N and LEW/Ss strains (* p < 0.05). The average load time is shown in panels **V-AB**. Tukey multiple comparison test detected several subtle differences between males and females at individual Sessions in the M520/N and F344/NCrl strains (* p < 0.05). All data shown as mean ± SEM.

**Table 4.** Linear regression analyses for males and females.

| Measure | Strain | Male | Female | Male v. Female F-test |
| --- | --- | --- | --- | --- |
| Acquisition: Infusions | M520/N | **$b = 0.70, p = 0.002^*$** | **$b = 1.47, p < 0.001^*$** | $F_{1,166} = 3.24, p = 0.074$ |
|  | WKY/NCrl | **$b = 1.27, p < 0.001^*$** | **$b = 0.57, p = 0.007^*$** | **$F_{1,136} = 4.65, p = 0.033^*$** |
| | F344/NCrl | **$b = 1.80, p < 0.001^*$** | **$b = 2.21, p < 0.001^*$** | $F_{1,166} = 0.57, p = 0.451$ |
| | F344/Stm | $b = 0.96, p = 0.164$ | **$b = 0.77, p = 0.007^*$** | $F_{1,146} = 0.04, p = 0.844$ |
| | LEW/Ss | **$b = 1.17, p = 0.028^*$** | $b = 0.20, p = 0.633$ | $F_{1,116} = 2.14, p = 0.146$ |
| | LEW/Crl | $b = 0.21, p = 0.706$ | $b = 0.40, p = 0.270$ | $F_{1,96} = 0.09, p = 0.770$ |
| | LE/Stm | **$b = 1.38, p < 0.001^*$** | **$b = 0.82, p = 0.003^*$** | $F_{1,116} = 1.58, p = 0.212$ |
| Acquisition: 15 min infusions | M520/N | **$b = 0.28, p = 0.005^*$** | **$b = 0.65, p < 0.001^*$** | **$F_{1,166} = 5.50, p = 0.020^*$** |
| | WKY/NCrl | **$b = 0.33, p < 0.001^*$** | **$b = 0.21, p = 0.001^*$** | $F_{1,136} = 1.29, p = 0.259$ |
|  | F344/NCrl | **$b = 0.28, p < 0.001^*$** | **$b = 0.56, p < 0.001^*$** | **$F_{1,166} = 5.29, p = 0.023^*$** |
| | F344/Stm | **$b = 0.45, p = 0.032^*$** | **$b = 0.22, p = 0.020^*$** | $F_{1,146} = 0.60, p = 0.439$ |
| | LEW/Ss | **$b = 0.40, p = 0.001^*$** | $b = 0.05, p = 0.637$ | **$F_{1,116} = 5.46, p = 0.021^*$** |
| | LEW/Crl | $b = 0.05, p = 0.695$ | $b = 0.16, p = 0.079$ | $F_{1,96} = 0.48, p = 0.492$ |
|  | LE/Stm | **$b = 0.57, p < 0.001^*$** | **$b = 0.23, p = 0.008^*$** | **$F_{1,116} = 5.32, p = 0.023^*$** |
| Acquisition: Latency to 1<sup>st</sup> infusion | M520/N | **$b = -0.39, p = 0.005^*$** | $b = -4.33, p < 0.001$ | **$F_{1,166} = 17.20, p < 0.001^*$** |
| | WKY/NCrl | $b = -1.39, p = 0.062$ | **$b = -2.66, p = 0.025^*$** | $F_{1,136} = 0.03, p = 0.872$ |
| | F344/NCrl | $b = -0.65, p = 0.427$ | **$b = -4.26, p = 0.016^*$** | $F_{1,166} = 3.83, p = 0.052$ |
| | F344/Stm | **$b = -4.13, p = 0.010^*$** | **$b = -6.37, p = 0.021^*$** | $F_{1,146} = 0.59, p = 0.443$ |
| | LEW/Ss | **$b = -1.74, p = 0.009^*$** | $b = -0.05, p = 0.929$ | **$F_{1,116} = 4.45, p = 0.037^*$** |
| | LEW/Crl | **$b = -0.16, p = 0.005^*$** | $b = -0.23, p = 0.063$ | $F_{1,96} = 0.27, p = 0.604$ |
| | LE/Stm | $b = -5.12, p = 0.017$ | $b = -1.24, p = 0.540$ | $F_{1,116} = 1.47, p = 0.227$ |
| Acquisition: Load time | M520/N | $b = -0.11, p = 0.361$ | **$b = -5.08, p < 0.001^*$** | **$F_{1,166} = 16.80, p < 0.001^*$** |
| | WKY/NCrl | **$b = -4.07, p = 0.004^*$** | $b = -3.61, p = 0.014$ | $F_{1,136} = 0.06, p = 0.815$ |
| | F344/NCrl | $b = -1.85, p = 0.133$ | $b = -7.05, p < 0.001$ | **$F_{1,166} = 6.70, p = 0.011^*$** |
| | F344/Stm | **$b = -3.96, p = 0.025^*$** | **$b = -7.74, p = 0.003^*$** | $F_{1,146} = 1.56, p = 0.214$ |
| | LEW/Ss | $b = -0.49, p = 0.215$ | $b = -0.19, p = 0.871$ | $F_{1,116} = 0.30, p = 0.582$ |
| | LEW/Crl | $b = -0.54, p = 0.285$ | $b = -0.45, p = 0.251$ | $F_{1,96} = 0.02, p = 0.882$ |
| | LE/Stm | **$b = -8.37, p = 0.001^*$** | $b = -4.61, p = 0.030$ | $F_{1,116} = 1.23, p = 0.270$ |
| Escalation: Infusions | M520/N | **$b = 6.13, p = 0.018^*$** | **$b = 9.07, p < 0.001^*$** | $F_{1,166} = 0.78, p = 0.379$ |
| | WKY/NCrl | $b = 6.93, p = 0.056$ | **$b = 4.635, p < 0.001^*$** | $F_{1,136} = 0.38, p = 0.537$ |
| | F344/NCrl | **$b = 4.35, p < 0.001^*$** | **$b = 8.81, p = 0.005^*$** | $F_{1,166} = 2.03, p = 0.156$ |
| | F344/Stm | $b = 3.70, p = 0.114$ | $b = 2.18, p = 0.316$ | $F_{1,116} = 0.18, p = 0.676$ |
| | LEW/Ss | $b = 3.05, p = 0.327$ | **$b = 11.83, p < 0.001^*$** | **$F_{1,116} = 4.28, p = 0.041^*$** |
| | LEW/Crl | $b = 4.73, p = 0.061$ | **$b = 7.88, p < 0.001^*$** | $F_{1,96} = 1.03, p = 0.312$ |
| | LE/Stm | **$b = 9.33, p < 0.001^*$** | **$b = 9.37, p < 0.001^*$** | $F_{1,116} = 0.00, p = 0.991$ |
| Escalation: Burst Episodes | M520/N | **$b = 0.87, p = 0.019^*$** | **$b = 1.77, p < 0.001^*$** | $F_{1,166} = 3.61, p = 0.059$ |
| | WKY/NCrl | $b = 0.23, p = 0.518$ | **$b = 0.48, p = 0.003^*$** | $F_{1,136} = 0.40, p = 0.526$ |
| | F344/NCrl | **$b = 0.59, p = 0.001^*$** | $b = 0.82, p = 0.054$ | $F_{1,166} = 0.27, p = 0.602$ |
| | F344/Stm | $b = 0.09, p = 0.656$ | **$b = 0.58, p = 0.010^*$** | $F_{1,146} = 2.44, p = 0.121$ |
| | LEW/Ss | $b = 0.10, p = 0.807$ | **$b = 1.08, p = 0.021^*$** | $F_{1,116} = 2.55, p = 0.113$ |
| | LEW/Crl | $b = 0.01, p = 0.981$ | $b = 0.58, p = 0.133$ | $F_{1,96} = 0.87, p = 0.353$ |
| | LE/Stm | **$b = 0.87, p = 0.002^*$** | $b = 0.44, p = 0.135$ | $F_{1,116} = 0.90, p = 0.346$ |

Significance indicated by emboldened text and asterisk (\*, $p < 0.05$ ).

Mixed-effect models were also used to determine if sex differences were present in the escalation of oxycodone intake **(Figure 11)**. Again, only significant main effects of Sex or the interaction between Sex and Session are reported. A significant main effect of Sex was observed for the WKY/NCrl strain in the total number of oxycodone infusions per session with males having higher intake compared with females (F_1, 12_ = 6.11, p = 0.029). A significant Sex-Session interaction was observed for the M520/N, F344/NCrl, and LEW/Ss strains with higher intake observed in female rats on several of the LgA sessions (M520/N: F_9, 135_ = 1.98, p = 0.047; F344/NCrl: F_9, 135_ = 2.94, p = 0.003; LEW/LEW/Ss: F_9,90_ = 2.03, p = 0.044). Regression modeling indicated robust positive slopes in both males and females in nearly all strains with female LEW/Ss rats displaying a significantly steeper slope than male LEW/Ss rats (**Figure 11**, **Table 4**). Using an unpaired t-test to compare the escalation index between males and females revealed no sex differences. Analysis of total oxycodone intake between males and females of each strain indicated higher intake in male WKY/NCrl rats compared with females (t_6.19_ = 2.47, p = 0.047), but no other strains. Analysis of the number of burst episodes during escalation revealed that female M520/N rats displayed significantly more burst episodes than male M520/N rats (F_1, 15_ = 10.01, p = 0.006). In contrast, male WKY/NCrl rats had a higher number of burst episodes than female WKY/NCrl rats that trended towards significance (F_1, 12_ = 3.99, p = 0.069). No differences between males and females were detected using regression modeling (**Table 4**). Considering that male M520/N rats displayed faster early session measures while female M520/N rats displayed a higher number of infusions and burst episodes during escalation, this suggests that distinct biological systems may govern the acquisition of oxycodone self-administration and dysregulation of oxycodone self-administration.

**Figure 11:**
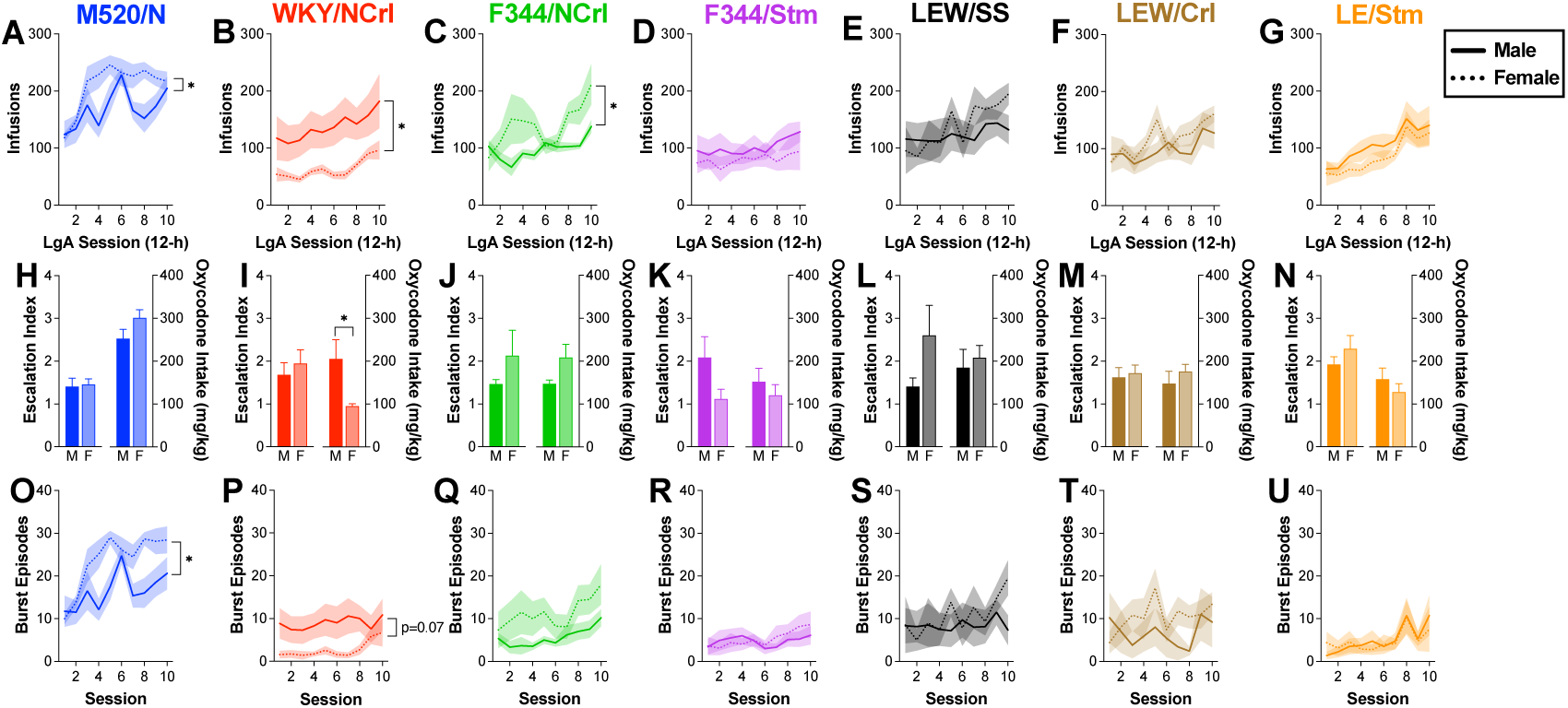
Comparison between male and females in escalation metrics. Total infusions per 12-h session during the escalation phase is displayed for males (solid line) and females (dashed line) in the M520/N (**A**), WKY/NCrl (**B**), F344/NCrl (**C**), F344/Stm (**D**), LEW/Ss (**E**), LEW/Crl (**F**), and LE/Stm (**G**) strains. M520/N and F344/NCrl females displayed greater oxycodone intake compared with M520/N and F344/NCrl females, while WKY/NCrl males displayed greater oxycodone intake compared with WKY/NCrl males (* p < 0.05). The escalation index (left) and total oxycodone intake (right) during the escalation phase is displayed for males (dark bar) and females (light bar) in panels **H-N**. No sex differences were observed in the escalation index for any strains. The WKY/NCrl males displayed significantly more total oxycodone intake compared with females (* p < 0.05). The number of burst episodes during each session is displayed for males (solid line) and females (dashed line) in panels **O-U**. Female M520/N rats displayed significantly greater burst responding compared with females (* p < 0.05) while male WKY/NCrl rats trended toward greater burst responding. All data shown as mean ± SEM.

## 4. Discussion

The use of inbred rat strains enables the identification of genetic contributions to specific OUD-related phenotypes that are difficult to detect in human studies. We analyzed a subset of inbred rat strains that displayed robust oxycodone intake escalation from our prior studies using a larger panel of inbred strains (21). We found that strains differed in the acquisition of oxycodone self-administration suggesting that genetic variability may influence the initial reinforcing effects of oxycodone. There were also significant strain differences in early session latency measures during acquisition supporting differences in the motivation and reinforcement learning associated with early self-administration of oxycodone. While all strains displayed an increase in oxycodone intake during the escalation phase, the M520/N strain self-administered significantly more oxycodone than all other tested strains. Moreover, the M520/N strain displayed a high rate of dysregulated oxycodone intake as measured by burst responding, a pattern of self-administration where several oxycodone self-infusions are delivered in a short period of time. These findings suggest that genetic factors contribute to distinct behavioral phenotypes, including within-session regulation of drug intake that may reflect the dysregulated and impaired control over oxycodone use.

In a study utilizing a similar IVSA paradigm, Doyle et al. (2023) examined burst responding in several inbred strains including the same WKY/NCrl and F344/NCrl strains assessed here. Consistent with our findings, they observed approximately 30% burst responding in the WKY/NCrl and F344/NCrl strains and similar within-session distribution of oxycodone infusions as measured by inter-infusion intervals. We demonstrate for the first time that the M520/N strain displays a striking pattern of oxycodone self-administration. In addition to showing relatively fast acquisition metrics, and high overall oxycodone intake during escalation, the M520/N strain engaged in significantly more burst responding with a significantly higher proportion of infusions (∼50%) occurring within bursts compared with other strains. Interestingly, female M520/N rats displayed a greater propensity to engage in burst responding compared with male M520/N rats suggesting that even within a genetically vulnerable strain, sex differences persist in the microstructure of drug-taking behavior and may reflect distinct underlying motivational or neurobiological mechanisms driving opioid intake.

The bursting pattern of intake displayed in some rats mirrors several features of human drug consumption that have been modeled in the intermittent access procedure. In this procedure, rats have short periods of drug access separated by longer intervals of abstinence (41). Intermittent access models are thought to capture cyclical patterns of intense drug-taking in humans (42). Some strains, particularly the M520/N strain, appear to naturally engage in a pattern of oxycodone use that is reflective of short bouts of intense drug-taking, making this a potentially clinically relevant phenotype. Although the limited access procedure was developed to mimic psychostimulant drug use, recent studies have incorporated intermittent access to heroin and oxycodone and found that it enhances drug craving throughout abstinence (28,43). Interestingly, burst-like responding to heroin self-administration was observed in both an intermittent-access model and a continuous-access model that removes the timeout period following heroin infusions in outbred Sprague Dawley rats (29). Burst responding is reminiscent of binge-like drug use, typically defined as intense drug consumption in a limited temporal window followed by a period of abstinence (44). Binge drug use is associated with increased risk for overdose (45). While binge patterns of opioid use are less reported in clinical studies, a survey of intravenous drug-users in Montreal found that binge use of opioids was observed in nearly 20% of all drug binge episodes with binge use of opioid/stimulant combinations occurring in approximately 14% of binge episodes (46). Although the specific temporal criteria for a binge differs depending on the drug class and pharmacokinetic profile, burst responding may reflect binge-like use during a single drug-taking session (39,47).

The inability to regulate drug intake through binge patterns of use has been associated with the development of addiction-related behaviors such as increased motivation, intake escalation, and relapse to drug seeking (37,38,48). For example, burst responding was associated with a higher preference for heroin compared to social reward in a drug-versus-social choice procedure in Sprague-Dawley rats (49). Other work in Sprague-Dawley rats demonstrated that temporal distribution of oxycodone infusions during escalation predicted subsequent reinstatement to oxycodone seeking in a sex-dependent manner. These findings suggest that the microstructural organization of intake may be a critical determinant of relapse risk (30). Together, these findings indicate that some strains, such as the M520/N strain, may provide useful genetic models for identifying the genes and biological mechanisms that promote dysregulated, binge-like opioid intake, and other potentially related phenotypes such as impulsivity, metabolism of oxycodone, and drug-induced plasticity that leads to OUD.

A recent study utilized an oral oxycodone self-administration paradigm to examine strain variation in oral oxycodone intake across 35 inbred rat strains (22). Interestingly, the M520/N strain displayed relatively low oral intake of oxycodone compared with the other strains tested. This contrasts with our observation that M520/N rats rapidly escalate intravenous oxycodone intake, display high burst responding, and have high overall oxycodone intake. It is plausible that oral and IVSA paradigms model partially overlapping but distinct components of drug reinforcement and intake patterns, and divergent intake patterns between the two models could be due to a variety of factors. For example, the metabolism and pharmacokinetics of oxycodone differ significantly between oral and intravenous procedures where potential first-pass elimination and delayed drug onset could impact its reinforcing effects when consumed orally (50,51). Oral procedures are also sensitive to taste aversions that may be more prevalent in some strains (52). Furthermore, the M520/N strain displayed high oral intake of alcohol compared with other inbred strains, even rat lines selectively bred for high alcohol consumption (53). Direct assessment of M520/N oxycodone metabolism across these routes of administration, as well as taste reactivity experiments, will shed light on the intake discrepancies observed between these procedures.

The Fischer (F344) and Lewis strains have been used as a genetic model of resistance and susceptibility respectively to substance use disorders, including OUD (54). Prior work found that the Lewis strain acquires heroin and morphine self-administration faster, display higher responding for both opioids under various opioid reinforcement schedules, and escalate intake at their preferred dose of heroin compared to the F344 strain (55–58). Mechanistically, strain differences in mu opioid receptor affinity and lower expression of endogenous opioids in the Lewis strain suggest a reduced basal tone of opioids (56,59). Comparisons of female LEW/Hsd and F344/Hsd rats in an oral oxycodone support the divergent intake patterns with LEW/Hsd females consuming more oxycodone, especially at higher oxycodone concentrations (23). More recent findings suggest more nuanced differences between the F344 and Lewis strains that is highly dependent on the substrains (22). We did not observe major differences in intravenous oxycodone intake between two F344 substrains (F344/NCrl and F344/Stm) and two Lewis substrains (LEW/Crl and LEW/SSNHsd). The Lewis and F344 substrains displayed similar oxycodone intake patterns with differences between the two strains in the early session metrics during acquisition where both Lewis substrains displayed significantly faster latencies to the first infusion and faster load times. This supports prior observations that the Lewis strain develops self-administration behavior more readily than the F344 strain. There were also subtle differences between the F344/NCrl and F344/Stm substrains in the inter-infusion intervals during the escalation phase. Differences between substrain behaviors can provide valuable insights into slight genetic divergences that could be useful in identifying genetic variants and establishing more refined genetic models (60–64). Together, these findings indicate that phenotypic differences between the Lewis and F344 strains are nuanced and substrain-dependent, underscoring the importance of genetic resolution when modeling vulnerability and resistance to opioid use disorder.

Male and female rats of each strain were compared using acquisition, escalation, and burst-related phenotypes. Analysis of the early session measures during the acquisition phase revealed sex differences in a subset of the tested strains. Of note, male M520/N rats displayed faster latencies to the first infusion and load times than female M520/N rats, while male F344/NCrl rats displayed faster load times compared with females. These differences were most apparent during the first few ShA acquisition sessions suggesting that males may engage in greater exploratory responding during the early portions of these sessions. Sex differences in the escalation measures were observed in the M520/N, WKY/NCrl, and F344/NCrl strains. Aligning with load time differences, male WKY/NCrl rats consumed significantly more oxycodone than female WKY/NCrl rats. In contrast, female M520/N and F344/NCrl rats displayed greater responding for oxycodone throughout the escalation phase. A recent study reported no differences between male and female oxycodone intake in the WKY/N and F344/N strains, despite males showing higher peak concentrations in plasma oxycodone (19). Prior work in the M520/N strain did not reveal sex differences in an oral oxycodone procedure, suggesting the differences observed in the present study may be specific to intravenous oxycodone self-administration (22). In the same study, female LE/Stm rats consumed more oral oxycodone compared with male LE/Stm rats, a sex-dependent effect that was not observed in the present study (22). In an earlier oral oxycodone study, it was reported that female LEW/Hsd rats consumed significantly more oxycodone that male LEW/Hsd rats (23). Although we did not see differences in overall oxycodone intake in either Lewis strain, the LEW/Ss females displayed a steeper slope during the escalation phase indicating a more rapid escalation of intake. Sex differences in the number of burst episodes during escalation sessions were only found in the M520/N strain, with females engaging in more burst responding than males. In a previous report, inter-infusion intervals and inter-burst intervals were found to predict reinstatement of responding following extinction of oxycodone self-administration behavior in males and females respectively, suggesting that burst responding in M520/N females may predict high reinstatement responding during abstinence (30). Together these findings suggest that sex differences in specific self-administration metrics were strongly strain-dependent, with males in some strains showing faster session parameters and females in other strains displaying greater intake escalation or burst responding patterns, highlighting that genetic background critically shapes sex-specific drug-taking profiles that are relevant to OUD risk.

In this study, we examined patterns of oxycodone use using between-session and within-session metrics, providing detailed phenotypes for each of the tested strains. Between-session measures can reveal differences in escalation of oxycodone use, while within-session measures provide insight into dysregulated patterns of use as well as motivation to obtain oxycodone. These distinct factors of substance misuse likely have a genetic basis, and self-administration paradigms utilizing inbred animal models can unveil these genetic contributions. As genetic screening becomes more common, identifying genetic backgrounds that are susceptible to escalation and dysregulated drug use will better inform clinicians about what populations may be susceptible to OUD. Further, while all strains presented in this study escalated their oxycodone use, identification of strains that are resistant to escalation may provide additional insights into populations that can safely utilize opioids with diminished risks for abuse. Opioids are effective analgesics and may pose significantly less risks if treatments are specifically tailored to individuals with a genetic resistance to oxycodone intake escalation.

## Supporting information

Supplemental Results

## Funding Statement

This study was supported by the National Institutes of Health, National Institute on Drug Abuse U01 DA051937 (Ehringer, Bachtell, Saba). Eamonn Duffy was supported by the National Institutes of Health, National Institute on Drug Abuse training grant (T32 DA017637). Laura Saba was supported by the National Institutes of Health, National Institute on Alcohol Abuse and Alcoholism R24 AA013162 (Tabakoff, Hoffman, Saba) and National Institutes of Health, National Institute on Drug Abuse P30 DA044223 (Williams, Saba).

## Data Availability

Data are available on request from the authors.

## Conflict of Interest

The authors declare that the research was conducted in the absence of any commercial or financial relationships that could be construed as a potential conflict of interest.

## Acknowledgements

The authors thank Melinda Dwinell PhD (Medical College of Wisconsin; R24OD024617) for providing several of the inbred strains studied in this article. We thank Safa Vaseemuddin and Tolulope Ajanaku for assistance with collecting behavioral data.

## Notes

### Competing Interest Statement

The authors have declared no competing interest.

