## Supplemental Results for "Genetic Modulation of Oxycodone Self-Administration Trajectories: From Initiation to Escalating Burst Patterns"

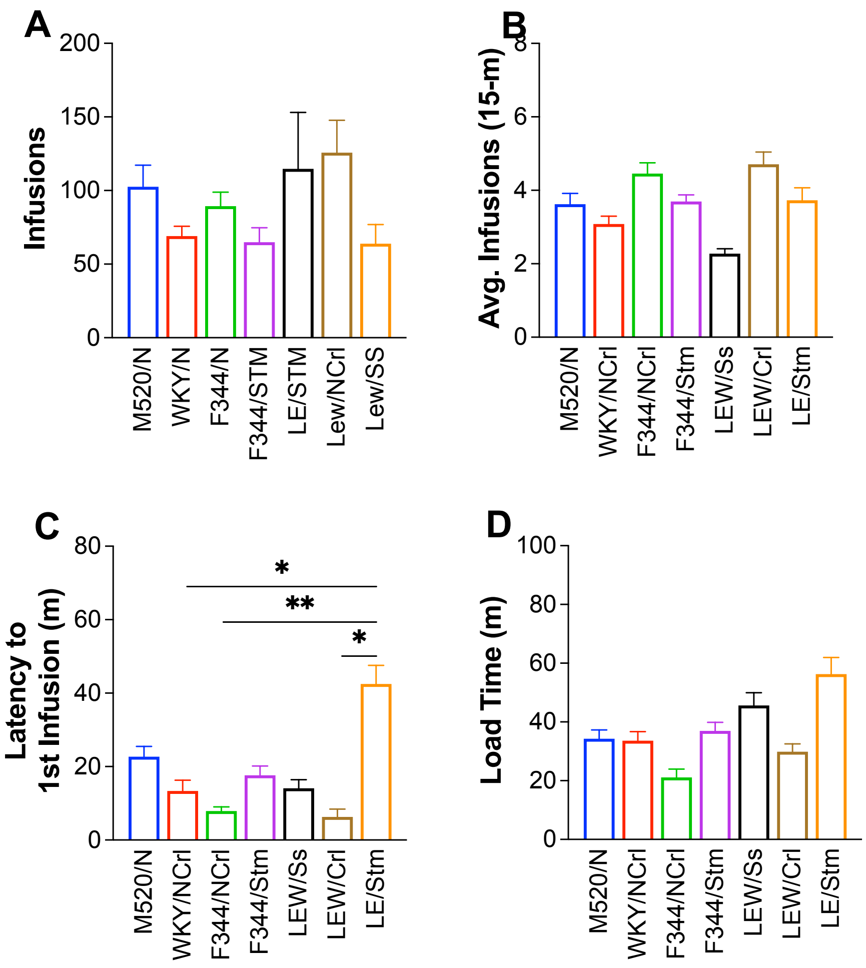


**Supplemental Figure 1: Acquisition Metrics for Saline Group.** (**A**) The total number of saline infusions during acquisition, (**B**) average number of infusions in the first 15-m of the session, (**C**) average latency to first infusion, and (**D**) average load latency during acquisition for the saline Group is displayed for the M520/N, WKY/NCrl, F344/NCrl, F344/Stm, LEW/Ss, LEW/Crl, and LE/Stm strains. A one-way ANOVA revealed no differences in total saline intake between strains. A one-way ANOVA compared the average latency to first infusion between strains and found significant differences between strains. Post hoc analysis found that the LE/Stm strain had a significantly longer latency to first infusion compared to the WKY/NCrl. F344/NCrl, and LEW/Crl strains ($p < 0.05$). A one-way ANOVA revealed no differences in average load times between strains. No strain differences were found in the average number of saline infusions during the first 15 minutes of a session. Significant differences between strains are highlighted with the symbol (* $p < 0.05$; ** $p < 0.01$).

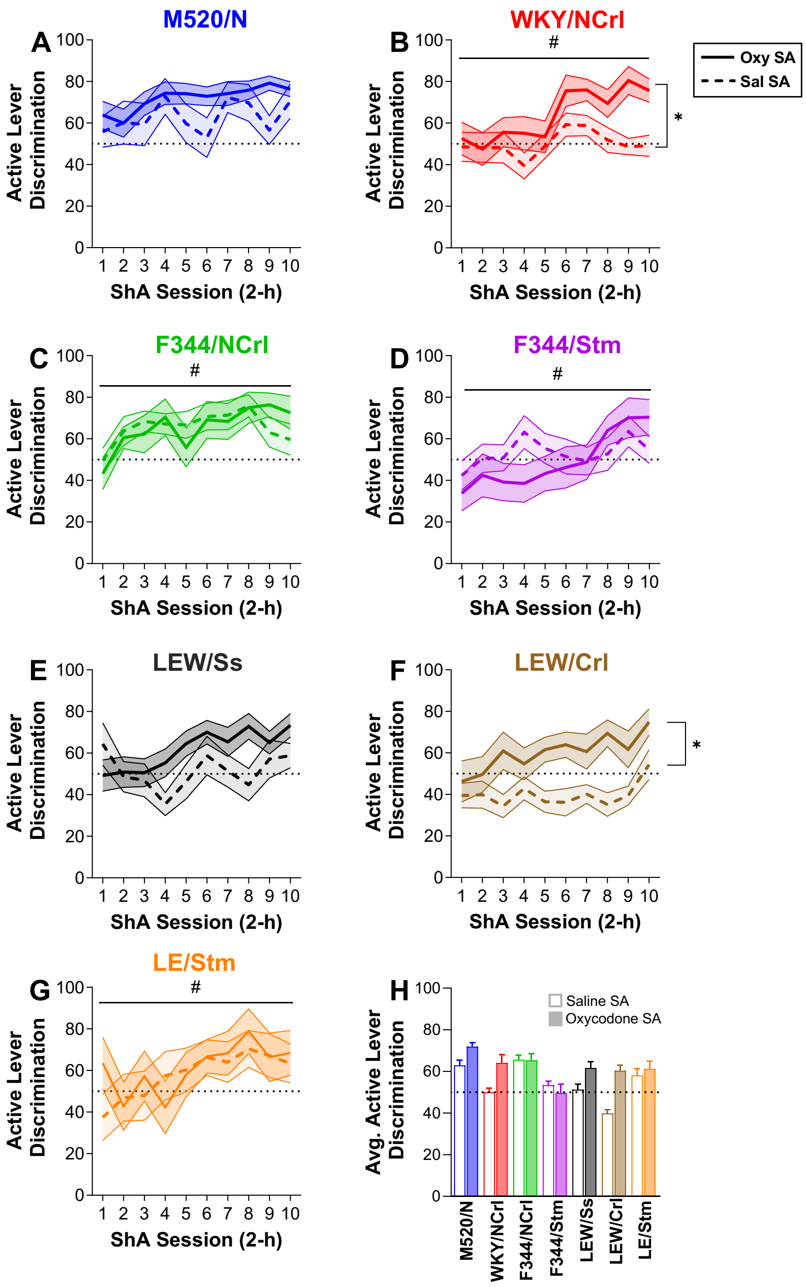


**Supplemental Figure 2: Lever discrimination during acquisition.** Lever discrimination across acquisition sessions is shown for oxycodone (solid line) and saline (dashed line) Groups in the M520/N (**A**), WKY/NCrl (**B**), F344/NCrl (**C**), F344/Stm (**D**), LEW/Ss (**E**), LEW/Crl (**F**), and LE/Stm (**G**) strains (mean ± SEM). WKY/NCrl and Lew/Crl showed main effects of Group (*, $p < 0.05$), while WKY/NCrl, F344/NCrl, F344/Stm, and LE/Stm showed main effects of Sesions (#, $p < 0.05$). (**H**) No significant differences were observed between strains were observed in the average active lever discrimination across the acquisition sessions in either the oxycodone or saline Groups. A horizontal dashed line is displayed on each line graph at 50% lever discrimination, with values above this threshold indicating a preference for the active lever.

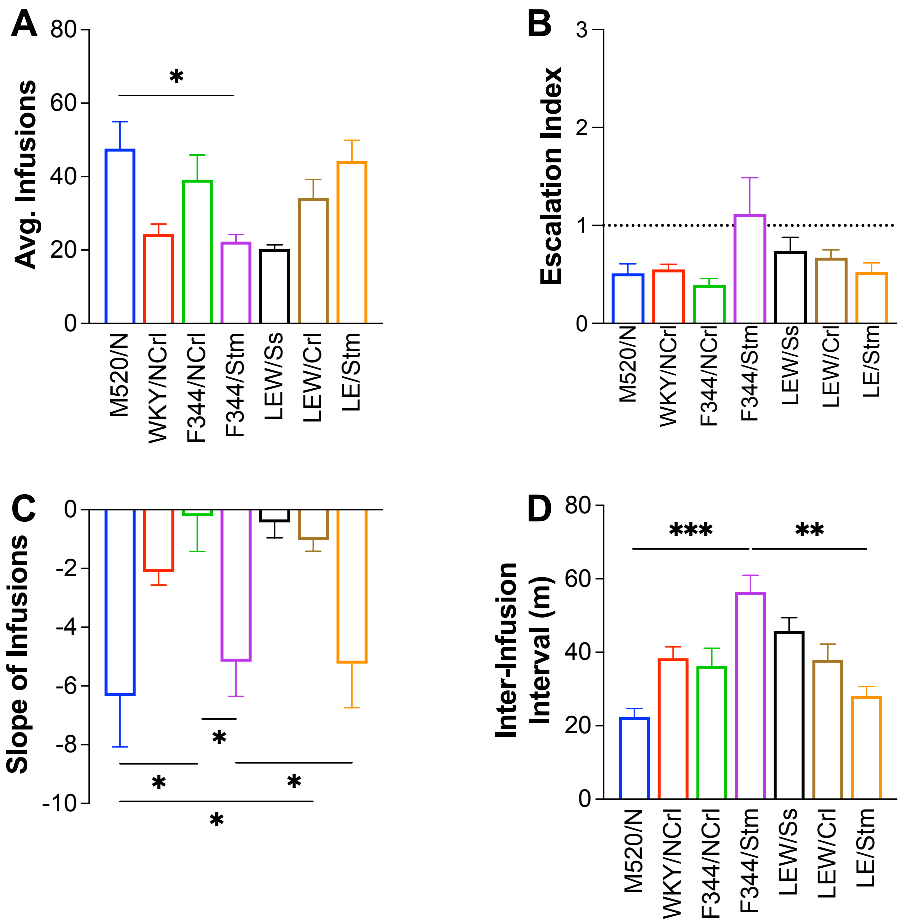


**Supplemental Figure 3: Escalation Metrics for Saline Group.** (**A**) The average number of saline infusions differed by strain with the M520/N rats displaying a higher number of saline infusions compared with the F344/Stm (* $p < 0.05$). (**B**) Escalation index was amongst the strains with most showing an escalation index below 1 indicating a decrease in saline intake across the escalation sessions. (**C**) Slope of infusions differed by strain with M520/N displaying a steeper negarive slope compared with Lew/Crl and F344/NCrl, and F344/Stm displaying a steeper negative slope compared with F344/NCrl and LE/Stm (* $p < 0.05$). (**D**) Average inter-infusion intervals also differed between the strains with F344/Stm rats showing the longest inter-infusion interval that was significantly different than M520/N and LE/Stm rats (** $p < 0.01$, *** $p < 0.001$). All data are plotted as mean ± SEM.

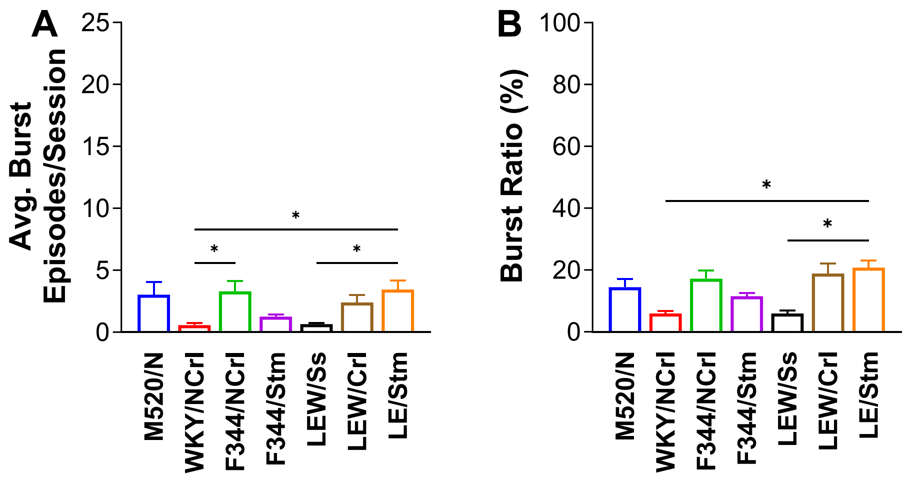


**Supplemental Figure 4: Burst Metrics in Saline Groups.** (**A**) Average number of bursts per session differed by strains with the WKY/NCrl displaying the fewest burst episodes. WKY/NCrl rats were significantly different that F344/NCrl and LE/Stm rats, and Lew/Ss rats were also significantly different than LE/Stm rats (* $p < 0.05$). (**B**) The burst ratio also differed between strains with LE/Stm rats displaying a higher proportion of saline infusions within bursts compared with Lew/Ss and WKY/NCrl rats (* $p < 0.05$). All data are plotted as mean ± SEM.
